# The N-terminal region of DNMT3A combines multiple chromatin reading motifs to guide recruitment

**DOI:** 10.1101/2023.10.29.564595

**Authors:** Hannah Wapenaar, Gillian Clifford, Willow Rolls, Hayden Burdett, Yujie Zhang, Gauri Deák, Juan Zou, Mark R. D. Taylor, Jacquie Mills, James A. Watson, Dhananjay Kumar, Alakta Das, Devisree Valsakumar, Janice Bramham, Philipp Voigt, Marcus D. Wilson

## Abstract

DNA methyltransferase 3A (DNMT3A) plays a critical role in establishing and maintaining DNA methylation patterns. However, the mechanisms underlying DNMT3A recruitment to and function within different chromatin environments remain unclear. Using a combination of biochemical and structural approaches we find that DNMT3A interacts using multiple interfaces with chromatin; directly binding generic nucleosome features as well as site-specific post-translational histone modifications. The N-terminal region, unique to the DNMT3A1 isoform, is essential for these interactions and stabilises H3K36me2-nucleosome recruitment. Intriguingly, in the same region critical for nucleosome binding we also map a ubiquitylation-dependent recruitment motif (UDR). The UDR binds specifically to ubiquitylated H2AK119, explaining the previously observed recruitment to Polycomb-occupied heterochromatin. A cryo-EM structure of DNMT3A1-DNMT3L with a modified nucleosome reveals that the UDR interacts with the nucleosome surface including the acidic patch. Previously unexplained disease-associated mutations are present in the UDR and ablate nucleosome interactions. This leads to an increased understanding of how DNMT3A1 recruitment occurs in the genome and highlights the importance of multivalent binding of DNMT3A to histone modifications and the nucleosome.

## Introduction

A pivotal player in the dynamic management of vertebrate genomes is DNA methylation, an epigenetic mark on cytosine bases which generally represses transcription. Upwards of 80% of CpG sites in the genome are methylated, but this is not equally distributed. Highly repetitive sequences are hypermethylated to ensure genome stability whereas promoters for developmental and housekeeping genes hypomethylated. Patterns of cytosine methylation are tissue-specific and heritable, which affects gene expression and subsequent cell fate determination. As a result, the correct maintenance and positioning of DNA methylation is critical to normal cellular function. Accordingly, aberrant DNA methylation patterns are found in numerous disorders including cancers, developmental and neurological diseases (Janssen & Lorincz, 2022; Plass *et al*, 2013).

DNA methyltransferases (DNMTs) catalyse the transfer of a methyl group from a S-adenosylmethionine (SAM) donor to the C5 position of a cytosine base (Song *et al*, 2012). Vertebrates have three methyltransferase proteins that deposit DNA methylation: DNMT1, DNMT3A and DNMT3B. DNMT3A and DNMT3B deposit *de novo* CpG methylation during differentiation, but show continued expression and activity into adulthood (Feng *et al*, 2005; Okano *et al*, 1999; Wu *et al*, 2010). Despite high conservation between the two proteins, DNMT3A and DNMT3B exhibit different chromatin binding preferences, different catalytic activity, and different interaction partners (Baubec *et al*, 2015; Gopalakrishnan *et al*, 2009; Morselli *et al*, 2015). There are two main isoforms of DNMT3A (Chen *et al*, 2003; Weisenberger *et al*, 2002): DNMT3A1 is the full-length form of the protein, whilst DNMT3A2 utilises an alternative promoter and as such lacks the divergent N-terminus of the full-length (Manzo *et al*, 2017). These isoforms show different expression patterns, with DNMT3A2 expressed during early development and in the germline and DNMT3A1 being the predominant postnatal isoform (Gu *et al*, 2022). However, the functional biochemical differences between isoforms have not been unravelled.

It is unclear how different regions of the genome recruit specific DNA methyltransferases. For instance, developmentally inactivated CpG islands are not normally methylated despite high CpG concentration (Brinkman *et al*, 2012). Also, DNA methyltransferases must contend with other chromatin proteins for access to substrate DNA. Most notably among these is the fundamental repeating unit of chromatin, the nucleosome. Comprised of two copies each of the four core histones, the nucleosome wraps and compacts ∼147 bp of DNA around its globular core (Luger *et al*, 1997). The nucleosome acts as a landing platform that integrates many signalling processes to help define genomic loci by extensive histone post-translational modification (Bannister & Kouzarides, 2011).

The recruitment of DNMT3A to specific genomic loci is a highly regulated process, finely tuned to ensure the fidelity of DNA methylation pattern. This is mediated in part, by direct recognition of multiple histone post-translational modifications. Beyond the C-terminal catalytic domain, DNMT3A contains chromatin recognition PWWP (Pro-Trp-Trp-Pro) (Dhayalan *et al*, 2010) and ADD (ATRX-DNMT3-DNMT3L) (Ooi *et al*, 2007) domains which help to target DNMT3A to non-transcribed intergenic regions marked by H3 Lysine 36 dimethylation (H3K36me2) and H3K4me0, respectively (Weinberg *et al*, 2019; Xu *et al*, 2020b). The PWWP domain directly recognises H3K36 methylation (Rondelet *et al*, 2016; Weinberg *et al*., 2019) and disruption of H3K36 methylation or binding leads to aberrant DNMT3A1 localisation and activity (Chen *et al*, 2022; Hamagami *et al*, 2023). However, the overall cellular and organismal phenotype is gain-of-function (Heyn *et al*, 2019; Sendzikaite *et al*, 2019), with aberrant hypermethylation at facultative heterochromatin Polycomb-associated regions. DNMT3A1 appears to localise to CpG island shores in both normal (Gu *et al*., 2022; Manzo *et al*., 2017) and disease state-mimic cells (Heyn *et al*., 2019; Sendzikaite *et al*., 2019; Weinberg *et al*, 2021). These regions are enriched with facultative heterochromatin marks H3K27me3 and H2AK119ub (Blackledge & Klose, 2021), but DNMT3A1 lacks recognisable domains for interaction with these marks. Removal of the DNMT3A N-terminal region affects recruitment (Gu *et al*., 2022; Weinberg *et al*., 2021). Exactly how DNMT3A integrates these multiple signals to ensure correct methylation at appropriate chromatin locations is unclear.

Here we biochemically map the N-terminal region and PWWP domain of DNMT3A1 as the minimal fragment required for robust binding to modified nucleosomes, particularly those marked with H3K36me2 or H2AK119ub. Disease-associated mutations in the N-terminal region were found to impair nucleosome binding, providing insights into the mechanistic basis of these mutations in various disease states. The same region implicated in binding the nucleosome surface also ensures specific interactions with H2AK119ub modified nucleosomes. Single-particle cryo-EM structures and subsequent analysis reveals DNMT3A1’s extensive engagement with the nucleosome acidic patch, H2A and H3, facilitating a highly specific interaction with H2AK119ub. Finally, despite its strong binding to nucleosomes, DNMT3A1’s catalytic activity was not proportional to its recruitment. These findings highlight a disconnect between DNMT3A recruitment and its enzymatic activity, shedding light on the complex relationship between DNMT3A and methylation of chromatinized DNA.

## Results

### The N-terminal region of DNMT3A1 facilitates interaction with chromatin

As DNA methylation occurs within the context of chromatin, we sought to understand which regions of DNMT3A are necessary for interaction on a minimal chromatinized nucleosome template. DNMT3A contains a C-terminal catalytic domain, highly similar to other DNA methyltransferases (Fig 1A). The N-terminal regulatory section comprises an ADD domain that interacts with the unmodified H3 tail (Ooi *et al*., 2007; Otani *et al*, 2009), a PWWP domain that binds H3 lysine-36 di-methylation (H3K36me2; Dhayalan *et al*., 2010; Rondelet *et al*., 2016; Weinberg *et al*., 2019) and a poorly characterised divergent N-terminal region. We expressed and purified different 6xHis-MBP tagged fragments of DNMT3A1’s regulatory section and reconstituted H3K36me2 nucleosomes (Figs S1A & S1B). Using electrophoretic mobility shift assays (EMSAs), we could observe robust binding of the DNMT3A regulatory section to modified nucleosomes *in vitro* (DNMT3A11^−614^; Fig 1B, S1C). The PWWP domain alone bound poorly, likely due to the low affinity of the interaction with the histone mark (DNMT3A1^278-427^; Dhayalan *et al*., 2010; Dukatz *et al*, 2019; Weinberg *et al*., 2019) and affinity for competitor DNA used in the assay buffer. Notably, removal any region after the PWWP domain, including the ADD domain had no measurable effect on interaction with H3K36me2-nucleosomes in our assay. This suggests that the minimal fragment required for nucleosome interaction was DNMT3A1^1-427^ comprising of the N-terminal region and PWWP domain (NT-PWWP; Fig 1B).

**Figure 1.**
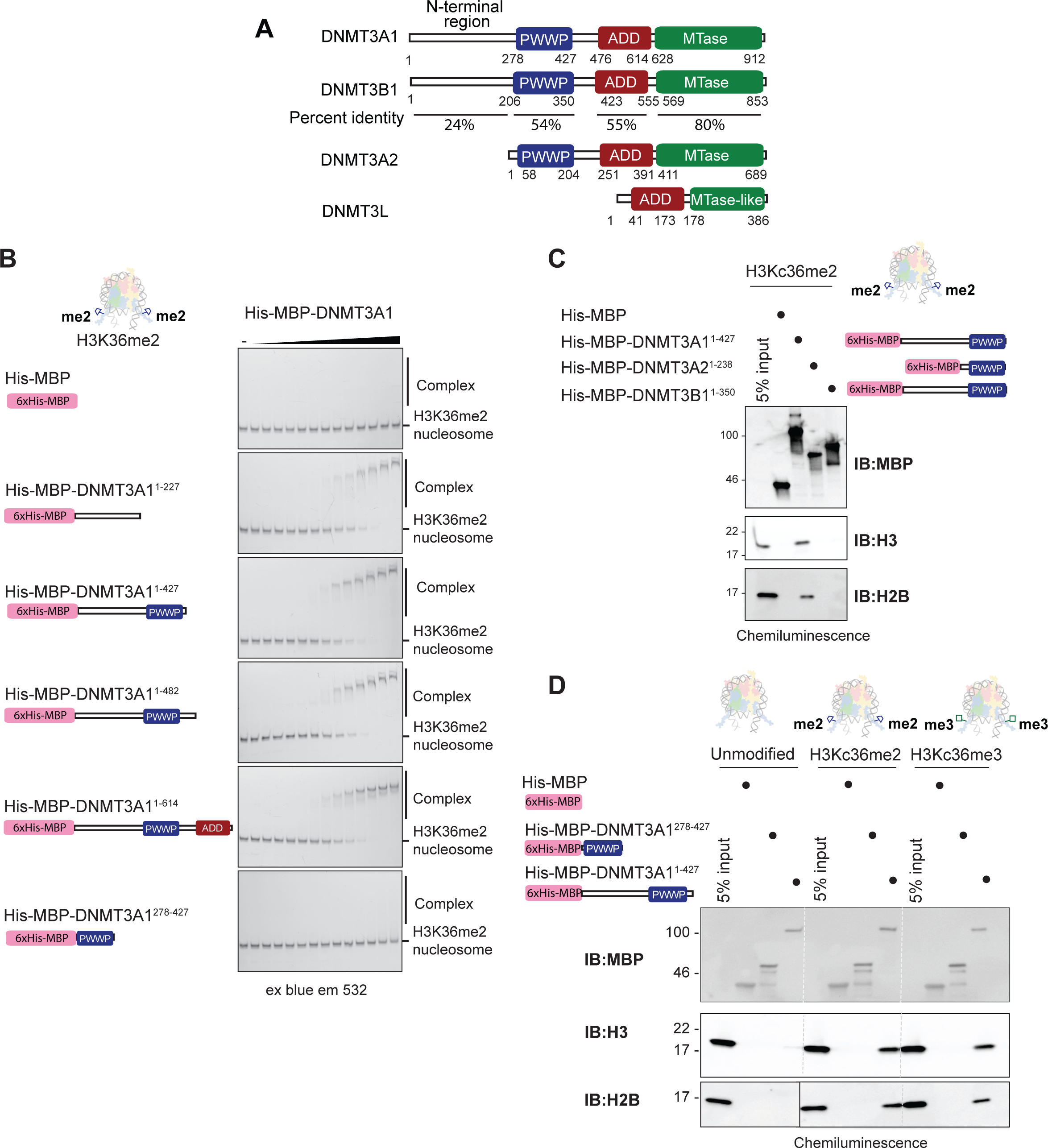
Mapping the minimal DNMT3A1 nucleosome-interacting fragment. A. Schematic domain overview of DNMT3A1, 3B1, 3A2 and 3L. PWWP = proline-tryptophan-tryptophan-proline domain, ADD = ATRX-DNMT3-DNMT3L domain, Mtase = methyl transferase domain. Percent amino acid identity per domain between DNMT3A1 and DNMT3B1 as determined by Uniprot sequence alignment is shown underneath relevant domains. B. Electrophoretic mobility shift assays (EMSAs) comparing interaction of different DNMT3A1 constructs with H3K36me2 nucleosomes wrapped with 5’ FAM labelled 175bp Widom601 DNA (FAM = fluorescein). Nucleosomes were incubated with increasing concentrations (0-8000 nM, 2x dilution series) of His-MBP-DNMT3A1 constructs. Complexes were resolved by native-PAGE and imaged using blue light excitation and 532nm emission filters. Experiment was repeated 2 times and quantification shown in Supp Fig S1C. C. Pull down assay comparing binding of 1 μg H3Kc36me2 nucleosomes to 12 μg amylose-bead immobilised DNMT3A1, 3A2 and 3B1 constructs from their N-terminus to the end of the PWWP domain. Bound nucleosomes were detected using western blot using against histones H3 and H2B. Anti-MBP antibody was used as loading control. D. Pull down assay comparing binding of DNMT3A1 (12 µg) constructs to unmodified, H3Kc36me2 and H3Kc36me3 nucleosomes (1 µg). Proteins were immobilised on amylose beads and bound nucleosomes were detected using western blot.

This result was particularly intriguing considering the N-terminal region has the highest degree of amino acid sequence divergence between DNMT3A and its paralog DNMT3B (Fig1A; Manzo *et al*., 2017). Furthermore, DNMT3A has two major isoforms which differ in this N-terminal region; the shorter DNMT3A2 isoform has been extensively studied biochemically and *in vivo* in mouse embryonic stem cells (Jeltsch & Jurkowska, 2016), but the predominant isoform found in somatic tissues is the longer DNMT3A1 that contains this N-terminal extension (Fig 1A; Chen *et al*, 2002; Gu *et al*., 2022; Manzo *et al*., 2017; Xu *et al*, 2020a). Indeed, using *in vitro* pull-down assays (Fig 1C) and EMSAs (Fig S1D) only the NT-PWWP region of DNMT3A1^1-427^ robustly interacted with modified nucleosomes compared to similar fragments from DNMT3B and DNMT3A2. This suggested that the DNMT3A1-specific feature within the N-terminal region could be a key point of difference between the *de novo* methyltransferases and may provide an explanation for tissue-specific DNA methylation patterns.

To investigate this possibility further we investigated binding to modified nucleosomes. The N-terminal region alone showed considerable interaction with nucleosomes irrespective of their modification state (Fig 1A), suggesting this region mediates direct nucleosome contacts. Nevertheless, interaction of DNMT3A1^1-427^ was further stimulated and specificity gained by recognition of H3K36me2/me3 (Fig 1D & S1E) in line with the observed localisation to H3K36-methylated regions of the genome (Dhayalan *et al*., 2010; Weinberg *et al*., 2019; Xu *et al*., 2020b). Furthermore, disrupting methylated-lysine recognition using Heyn-Sproul-Jackson syndrome causing mutations in the PWWP domain (Heyn *et al*., 2019) removed H3K36-methylation specificity (Fig S1E). Overall, this data shows the unique DNMT3A1 NT-PWWP region is the minimal chromatin-interacting fragment required for robust H3K36me2-nucleosome recognition.

### DNMT3A1 N-terminal region contains DNA and nucleosome binding regions

To precisely map the region of the DNMT3A1^1-427^ NT-PWWP fragment that is important for nucleosome interaction, we reacted zero-length covalent crosslinkers (see Methods) with preformed DNMT3A1^1-427^:H3Kc36me2-nucleosome complex (Fig S2A) and identified crosslinked residues by mass spectrometry (Fig 2A). Multiple repeat experiments revealed similar crosslinking patterns, including validating crosslinks between histones as predicted by the nucleosome structure (Ai *et al*, 2022; Luger *et al*., 1997). Several crosslinks between the N-terminal tail of H3 and PWWP domain we observed, as expected given the H3K36me2 interaction. Intriguingly, the most predominant crosslinks clustered between residues 120-200 of DNMT3A1 and H2A-H2B (Fig 2A & S2B). The crosslinks observed between H2A and H2B were either part of or adjacent to the nucleosome acidic patch (Fig 2B), which is a negative charged recess on the nucleosome surface and a common site of interaction for chromatin binding proteins (McGinty & Tan, 2021). Neutralising the charge on the acidic patch through mutation (*acidic patch*: H2A^E61A/E91A/E92A^ and H2B^E105A^) greatly reduced binding of DNMT3A1^1-427^ to both H3K36me2 nucleosome (Fig 2C) and unmodified mutant nucleosomes (Fig S2C), suggesting the acidic patch region is required for DNMT3A1 nucleosome association.

**Figure 2.**
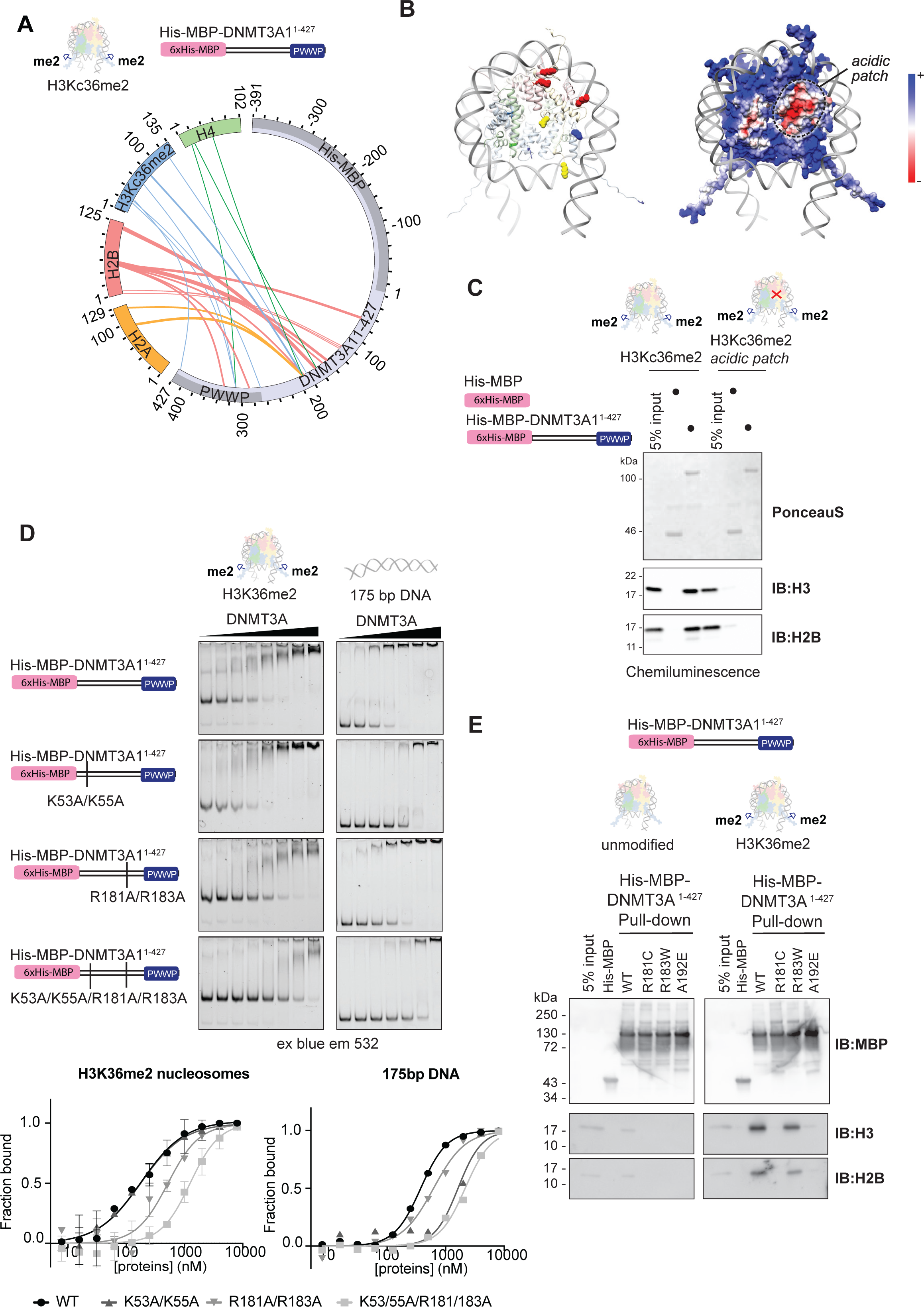
DNMT3A1 forms discreet contacts with the nucleosome surface and DNA. A. Crosslinking mass spectrometry of His-MBP-DNMT3A1^1-427^ to H3Kc36me2 nucleosomes wrapped with 175bp Widom-601 DNA. Circular representation shows two biological replicates combined of which one measured in triplicate. Only crosslinks between DNMT3A1 and histones are shown. Crosslinks weighted based on the number of times the high-confidence crosslink appears in the 4 replicates (thin = 1x, medium = 2x, wide = 3x). B. Left: Crosslinks from Fig 2A mapped on a nucleosome model (PDB 1AOI (Luger *et al*., 1997)). More abundant crosslinked residues shown as spheres, less as stick representation. Model coloured, Yellow, H2A; Red, H2B; Blue, H3; Green, H4. Right: nucleosome model in same orientation as left coloured based on surface charge, to highlight the acidic patch region (dashed circle). Blue indicates positive charge, red indicates negative charge. C. Pull down assay comparing DNMT3A1 binding to H3Kc36me2 nucleosomes with or without mutations in the acidic patch (H2A^E61A/E91A/E92A^ and H2B^E105A^). DNMT3A1^1-427^ or tag alone was immobilised on amylose beads and proteins were detected using PonceauS stain. Bound nucleosomes were detected using western blotting for histones H3 and H2B. D. EMSA comparing binding of DNMT3A1 with mutations in the N-terminus to H3K36me2 nucleosomes or free 5’ FAM labelled 175bp DNA. 5’ FAM labelled tracer was incubated with 0-8000 nM (2x dilution series) of His-MBP-DNMT3A1^1-427^ variants. Gels show concentrations 62.5-8000 nM for nucleosomes and 125-8000 nM for free DNA, quantification was done with full concentration series. Experiment was repeated once (DNA) or two times and quantified (nucleosome). E. Pull down assay to investigate the effect of DNMT3A1 N-terminal region missense mutations in found in clinical patients on binding to H3K36me2 nucleosomes. Equal amounts of wild type and mutant His-MBP-DNMT3A1^1-427^ was immobilised on amylose beads and incubated with nucleosomes prior to washing and detection by western blot.

We next sought to biochemically map the DNMT3A1 interaction with the nucleosome acidic patch and other features further. We purified fragments of DNMT3A1 N-terminal-PWWP region lacking stepped-intervals of ∼50 residues from the N-terminus (Fig S1A). In agreement with the crosslinking mass spectrometry findings, the greatest loss of binding was observed between DNMT3A1^142-427^ and DNMT3A1^192-427^ (Fig S2D), mapping the acidic patch interaction within this region of DNMT3A1. A previous study identified two separable N-terminal basic clusters implicated in DNMT3A1 DNA binding (Suetake *et al*, 2011). In agreement, removal of the positive charge of the first region at Lys-53 and Lys-55 in DNMT3A1^1-427^ greatly reduced interaction to a 175bp DNA fragment but not H3K36me2-nucleosome (Fig 2D & S2E). In contrast, R181A and R183A mutations ablated the nucleosome interaction, but only marginally affected DNA binding (Fig 2D & S2F). As Arg-181 and Arg-183 are adjacent to the highly cross-linkable region and are absent in the most severe deletion fragment observed, we hypothesised that rather than DNA binding, these residues interact with the nucleosome acidic patch. Mutation of all 4 residues (K53A/K55A/R181/R183A) reduced overall DNA binding similarly to the K35A/K55A. However, this was additive to the R181A/R183A mutation on H3K36me2-nucleosomes, suggesting Lys-53/55 can contact nucleosome-linker DNA contributing to overall nucleosome interaction.

Disease-associated mutations of Arg-181 and Arg-183 have been reported in clonal haematopoiesis (Jaiswal *et al*, 2014), Tatton-Brown-Rahman syndrome (Tatton-Brown *et al*, 2018) and cancer patients (Basturk *et al*, 2017; Dutton-Regester *et al*, 2013; Huang *et al*, 2022), with no reported mechanism. These mutations do not affect protein stability but reduce DNMT3A1’s capacity to repress transcription in a reporter assay (Huang *et al*., 2022). We found R181C, and to a lesser extent R183W, reduced overall binding to H3K36me2 nucleosomes (Fig 2E), suggesting a mechanistic basis for the role of these mutations in disease states. Overall, the minimal NT-PWWP fragment of DNMT3A1 incorporates recognition of generic chromatin features to increase overall affinity to chromatin, with nucleosome surface recognition mapped to a central region of the N-terminal region.

### H3K36me2 stimulates DNMT3A1-DNMT3L complex activity on nucleosomes

To test whether our observations on shorter fragments of DNMT3A1 hold true for larger assemblies of DNMT3A1, we co-expressed full-length DNMT3A1 with its binding partner DNMT3L in *E. coli* and purified the complex to near homogeneity (Fig 3A,B & S3A,B). DNMT3A1-DNMT3L was soluble (Fig S3B) and found in two populations, possibly indicative of a heterotetrametric and heterodimeric states (Holz-Schietinger *et al*, 2011; Jia *et al*, 2007; Zeng *et al*, 2020). To test DNMT3A1-DNMT3L activity we used the strong positioning Widom-601 DNA sequence flanked by a single 50 bp linker region as a substrate. This was engineered to contain dual CpG targets optimised for DNMT3A binding and activity (Fig S3E; Gao *et al*, 2020; Mallona *et al*, 2021). We then assayed turnover of the methyl donor cofactor S-adenosylmethionine (SAM, Fig 3C & S3C,D). Activity was poor on DNA alone but substantially stimulated when the same sequence was used on wrapped nucleosomes (Fig 3B & Fig S3F). This affect is likely due to release of the normally autoinhibitory domain of the ADD, upon binding to the unmodified H3 tail (Guo *et al*, 2015). Indeed, nucleosomes lacking the first 24 residues of H3 in the normally flexible tail (termed ‘H3 tailless’) did not lead to significant increased activity compared to DNA alone (Fig 3C). Methylation of H3K36 in nucleosome substrate stimulated the activity of DNMT3A1-DNMT3L. This elevated activity is presumably mediated through increased recruitment of the enzyme to substrate but may also be due to release of PWWP mediated inhibition and/or increased availability of substrate methylation sites (Brohm *et al*, 2022).

**Figure 3.**
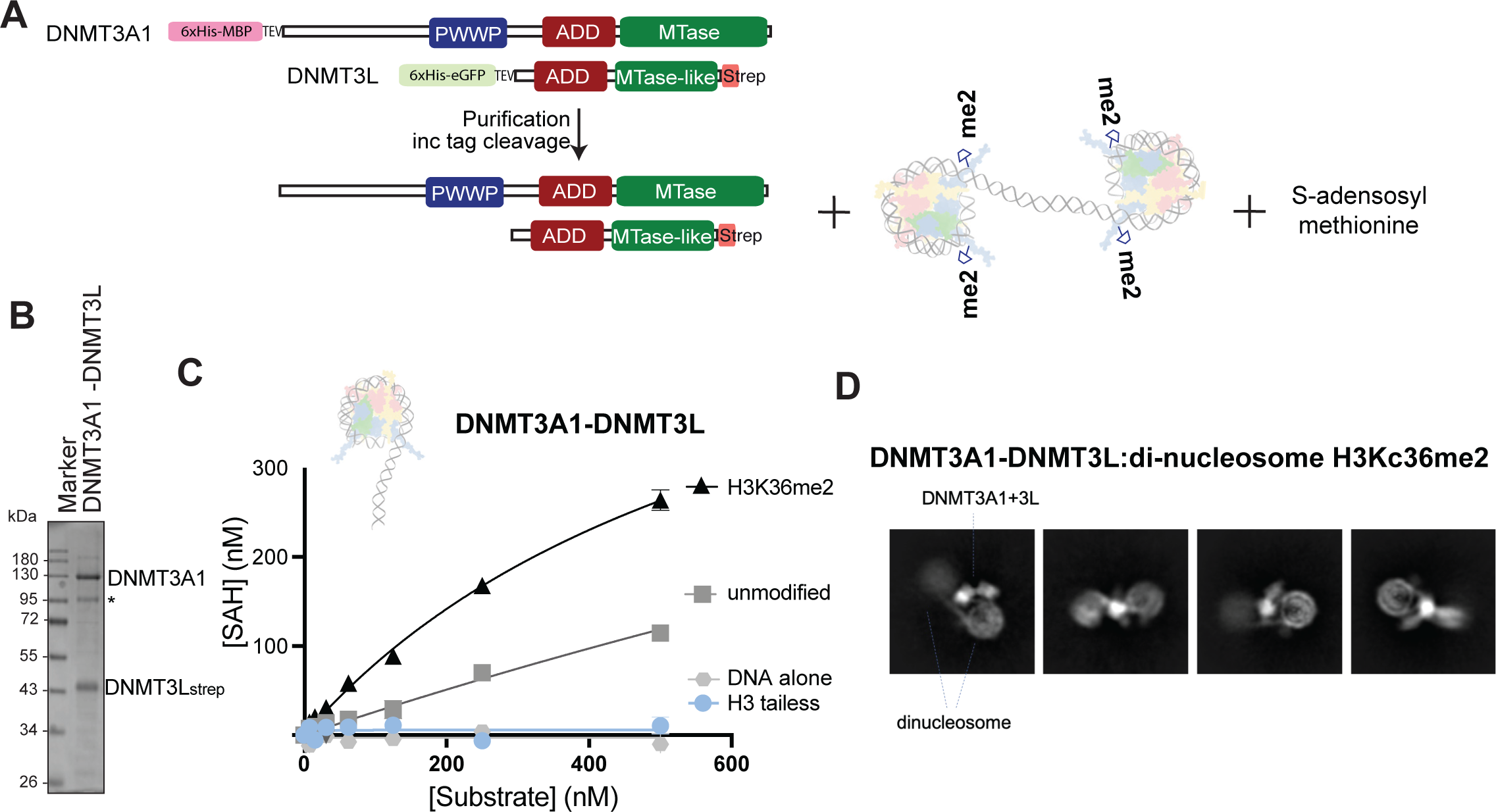
Full-length DNMT3A1-DNMT3L complex can methylate nucleosomal DNA and interacts with di-nucleosomes. A. Schematic overview of constructs used for cryo-electron microscopy and enzymology experiments. Tagged Full-length DNMT3A1 and DNMT3L were copurified and their tags cleaved. For cryo-EM they were incubated with SAM cofactor and di-nucleosome template. B. SDS-PAGE gel of purified full-length DNMT3A1-DNMT3L-StrepII complex. Asterisk indicates degradation product. C. Methyltransferase assay of full-length DNMT3A1-DNMT3L-StrepII on unmodified, H3K36me2, H3 tailless (H3 Δ1-24 A25C) nucleosomes wrapped with 195 bp Widom-601 DNA and free 195 bp Widom-601 DNA. DNMT3A1-DNMT3L was incubated with increasing concentrations of nucleosome/DNA substrate for 1 hour at 37°C. Methyltransferase activity was detected using Promega MTase-Glo™ Methyltransferase Assay. Michaelis-Menten curves were fit using GraphPad Prism 10. Experiment performed in duplicate. D. Selected 2D class averages obtained for DNMT3A1-3L: di-nucleosome H3Kc36me2 sample obtained during image processing.

To further understand how DNMT3A1-DNMT3L interacts on chromatin substrate we reconstituted the methyltransferase complex with H3Kc36me2 modified di-nucleosomes (Fig S3G & H). Given the wealth of chromatin binding regions in the proposed complex we surmised that di-nucleosomes would provide a better representation of the multivalent interactions expected of this complex. Using single-particle cryo-EM with full-length DNMT3A1-DNMT3L-StrepII complexed with di-nucleosomes in the presence of the cofactor SAM (Fig S3A), we could observe individual di-nucleosome particles (Fig S3I) and these averaged to reveal characteristic nucleosome-like 2D projections (Fig 3D). One of the nucleosomes was less well ordered than the other suggesting a degree of flexibility across linker DNA between the two protomers. Interestingly, we could also observe some additional density ascribed to DNMT3A1-DNMT3L on linker DNA but closer to the better ordered nucleosome-protomer. Due to severe preferred sample orientation, we were unable to reliably determine a 3D reconstruction from this data. Nevertheless, our experiments show that nucleosome substrates are preferentially methylated and that DNMT3A1-DNMT3L can engage linker DNA.

### The N-terminal region of DNMT3A1 binds to H2A Lys119ub within a nucleosome

Having mapped interaction with H3K36me2 modified nucleosomes we wanted to explore interactions with other DNMT3A1-bound chromatin states. There appear to be two competing mechanisms for DNMT3A recruitment *in vivo*, with enrichment at intragenic regions marked with histone H3K36me2 but also tissue-specific recruitment to H3K27me3-H2AK119ub marked facultative heterochromatin (Gu *et al*., 2022; Heyn *et al*., 2019; Manzo *et al*., 2017; Sendzikaite *et al*., 2019; Weinberg *et al*., 2021). It has recently been shown that direct recruitment to Polycomb regions is mediated by the N-terminal region of DNMT3A1 and was bioinformatically predicted to map to the same region that we determined to be important for nucleosome interaction (Gu *et al*., 2022; Weinberg *et al*., 2021). To investigate if this recruitment is direct, we generated nucleosomes marked with H3K27me3. In line with previous findings (Gu *et al*., 2022; Weinberg *et al*., 2019; Weinberg *et al*., 2021), H3K27me3 does not promote DNMT3A1^1-427^ interaction with nucleosomes (Fig S4A), nor stimulates enzymatic activity on nucleosomes (Fig S4B).

H3 Lys27 di-methylation (H3K27me2) is also deposited by Polycomb complex PRC2, but unlike H3K27me3 is widespread throughout the genome co-occurring with H3K36me2 (Ferrari *et al*, 2014; Mao *et al*, 2015; Streubel *et al*, 2018). Nucleosomes bearing both H3 di-methylation marks bound similarly to just H3K36me2 marked nucleosomes (Fig 4A). Taken as a whole, this points to PRC2 mediated H3K27 methylation neither directly inhibiting nor stimulating DNMT3A1 interaction. However, H3K27me3 does guide the deposition of the other Polycomb mark H2A Lys-119 ubiquitylation (H2AK119ub). To test the hypothesis that H2AK119ub directly targets DNMT3A1 to Polycomb regions we created ubiquitylated histones. Using a chemical alkylation approach, we cross-linked ubiquitin to position 119 of H2A, purifying the H2A-ubquitiylated mimic (H2AKc119ub; Long *et al*, 2014; Wilson *et al*, 2016; Burdett *et al*, 2023) and reformed these into nucleosomes (Fig S4C). Remarkably, in contrast to H3K27 methylation, H2AKc119ub nucleosomes bound with high affinity to DNMT3A1^1-427^ (Fig 4A). This suggests that the observed DNMT3A1 recruitment to Polycomb regions is mediated via direct binding to PRC1-mediated H2A ubiquitylation rather than H3K37me3 interaction, in line with recent findings (Gu *et al*., 2022; Weinberg *et al*., 2021).

**Figure 4.**
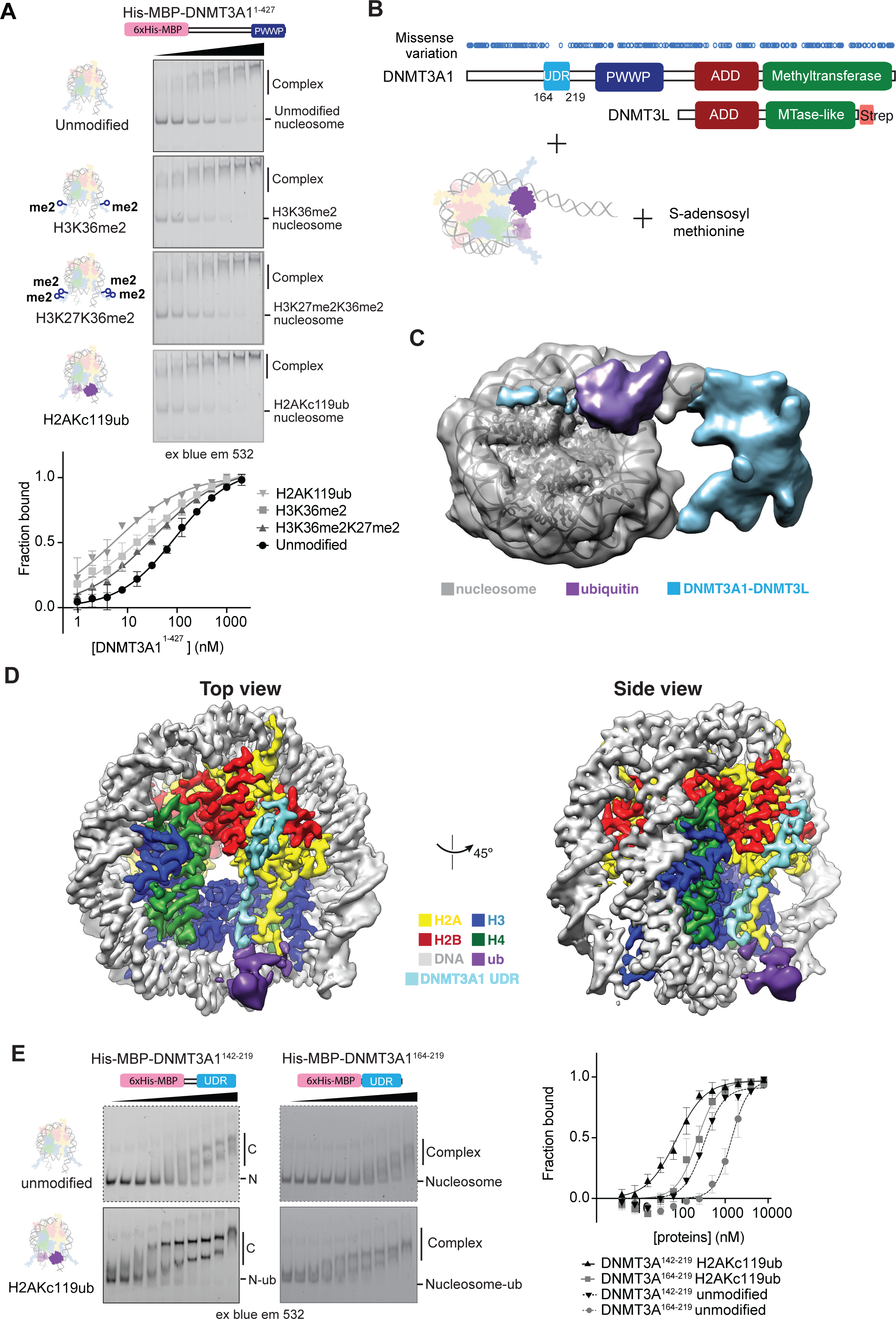
DNMT3A1 binds to H2AK119ub marked nucleosomes using a N-terminal UDR. A. EMSA comparing binding of DNMT3A1^1-427^ to unmodified, H3K36me2, H3K27me2K36me2 and H2AKc119ub nucleosomes. nucleosomes were incubated with increasing concentrations (0-2000 nM, 1.5x dilution series) of His-MBP-DNMT3A1^1-427^. Gels show concentrations 175nM-2000 nM, quantification was done with full concentration series, from an experiment in duplicate. B. Schematic overview of constructs used for cryo-electron microscopy. DNMT3A1-DNMT3L-StrepII was incubated with H2AK119ub nucleosomes wrapped with 195bp DNA and SAM cofactor. 1D plot of known missense mutations (blue ovals) taken from the gnomAD database, shows region with large sparsity of missense mutations in population (0.26 = frequency of variants in domain/frequency of variants in full length protein). C. Surface rendering of the 5.1 Å DNMT3A1-3L: H2AK119ub Nucleosome. Locally filtered segmented map with nucleosome density displayed at 0.006 threshold and ubiquitin and UDR density displayed 0.05. Rigid body fitting of nucleosome model (PDB 7VVU (Qu *et al*, 2022)), guided segmentation to identify non-nucleosome features including ubiquitin/UDR density (purple), nucleosome contacting UDR density (cyan) and linker DNA adjacent DNMT3A1-DNMt3L density (cyan). D. Surface rendering of the local resolution-filtered focused 3.1 Å resolution DNMT3A-3L: H2AK119ub Nucleosome complex map viewed along the DNA axis and the 45° rotation. Density was segmented and coloured according to local histone (coloured by convention), DNA (grey), Ubiquitin (purple) and UDR (cyan) features. Ubiquitin could not be readily placed in the attributed density and is displayed at 0.13 threshold, compared to the rest of the map at 0.35. E. EMSA comparing binding of the UDR of DNMT3A1 (DNMT3A1^164-219^) or extended UDR (DNMT3A1^142-219^) to unmodified and H2AKc119ub nucleosomes. nucleosomes were incubated with increasing concentrations (0-8000 nM, 2x dilution series) of His-MBP-DNMT3A1 fragments and resolved by native-PAGE. Imaging for Fluorescent DNA signal was performed using blue light excitation and 532nm emission filters. Gels show concentrations 15.6-8000 nM, quantification was done with full concentration series from two experiments.

### Structure of DNMT3A1-DNMT3L bound to a H2AKc119ub-nucleosome reveals DNMT3A1 interactions on the nucleosome surface

To understand how DNMT3A1 can specifically interact with ubiquitylated H2A on a nucleosome, we wrapped H2AKc119ub nucleosomes with 195 bp DNA and preformed a complex with DNMT3A1-DNMT3L-StrepII in the presence of SAM cofactor (Fig 4B) and determined the structure by single particle cryo-EM (Fig S5). Individual nucleosome-shaped particles could be observed in the raw data and 2D class averages were reminiscent of nucleosome structures, with some additional density attributable to ubiquitin and DNMT3A1 (Fig S6A & B). From our 3D reconstruction the assembly appeared to be highly mobile, with abundant flexible density on the linker DNA attributed to the catalytic core of the DNMT3A1-DNMT3L complex (Fig 4C), limiting the global resolution to 5.1 Å (Fig S5 Map 1, Fig S6C &D). We presume this density is weaker due to linker DNA flexibility and multiple conformations of the DNMT3A1-3L complex on DNA. The nucleosome core displayed higher resolution (Fig S5) with clear density for DNA and protein components of the nucleosome with some extra density on the nucleosome surface and adjacent to exit DNA. Focussed masking of only the better resolved nucleosome and adjoining densities allowed us to determine a structure at 3.1 Å resolution overall (Fig 4D, Fig S5 Map 2-4 & S6E-G). In this map, the histone core of the nucleosome and bound DNA are well ordered and similar to other nucleosome structures (RMSD 0.39 Å, PDB 7XD1; Ai *et al*., 2022), with the extended linker DNA projecting away from the nucleosome core. We could attribute density over the C-terminal tail of H2A to ubiquitin, which appears better ordered on one face of the symmetrical histone octamer compared to the other, co-incident with the linker DNA and DNMT3A1-DNMT3L densities. On the nucleosome surface, continuous DNMT3A1 density snakes in a N-C direction from the acidic patch region to become sandwiched in a depression formed between H3’-H3 and H2A prior to interaction with ubiquitin tethered over the C-terminal tail of H2A. We were able to build a discontinuous model of coiled structure into the density for DNMT3A1 corresponding to residues 166-171 and 177-194, with a buried surface interface of 2595 Å^2^ (Fig S5H).

To validate our model and further map the minimal DNMT3A1 fragment required for interaction with nucleosome-H2AK119ub, we sequentially deleted the N-terminal region. This confirmed that the same residues within 142-192 are required for H2AK119ub- and H3K36me2-nucleosome interaction (Fig S4D), albeit with a more radical effect for DNMT3A1^192-427^. In isolation a region comprising residues 142-219 is sufficient for preferential interaction with H2AKc119ub nucleosomes (Fig 4E, FigS4E). We further mapped this region and retained ubiquitin selectivity and affinity from DNMT3A1^164-219^, although with overall lower binding affinity to both unmodified and ubiquitylated nucleosomes (Fig 4E). Overall, a smaller fragment is required for H2K119ub binding in contrast to H3K36 methylation recognition. The PWWP domain is dispensable for ubiquitylated-nucleosome interactions, explaining why PWWP-histone-interaction mutations do not affect recruitment to Polycomb regions.

It was hypothesised that the N-terminal region contains a ubiquitylation-dependent recruitment motif (UDR; Weinberg *et al*., 2021) and we show that this maps to 164-219 of DNMT3A1, integrating both ubiquitin recognition as well as our nucleosome interaction region (Fig 5A). Intriguingly, this region is found to be highly invariant in the normal population, based on the absence of missense variants found in the gnomAD population database (Fig 4B; Deak & Cook, 2022; Karczewski *et al*, 2020). This underlines the importance of the DNMT3A1 nucleosome and ubiquitin interacting region during normal development and agrees with a recent mouse study showing the necessity of the N-terminal region of DNMT3A1 for postnatal neural development (Gu *et al*., 2022).

**Figure 5.**
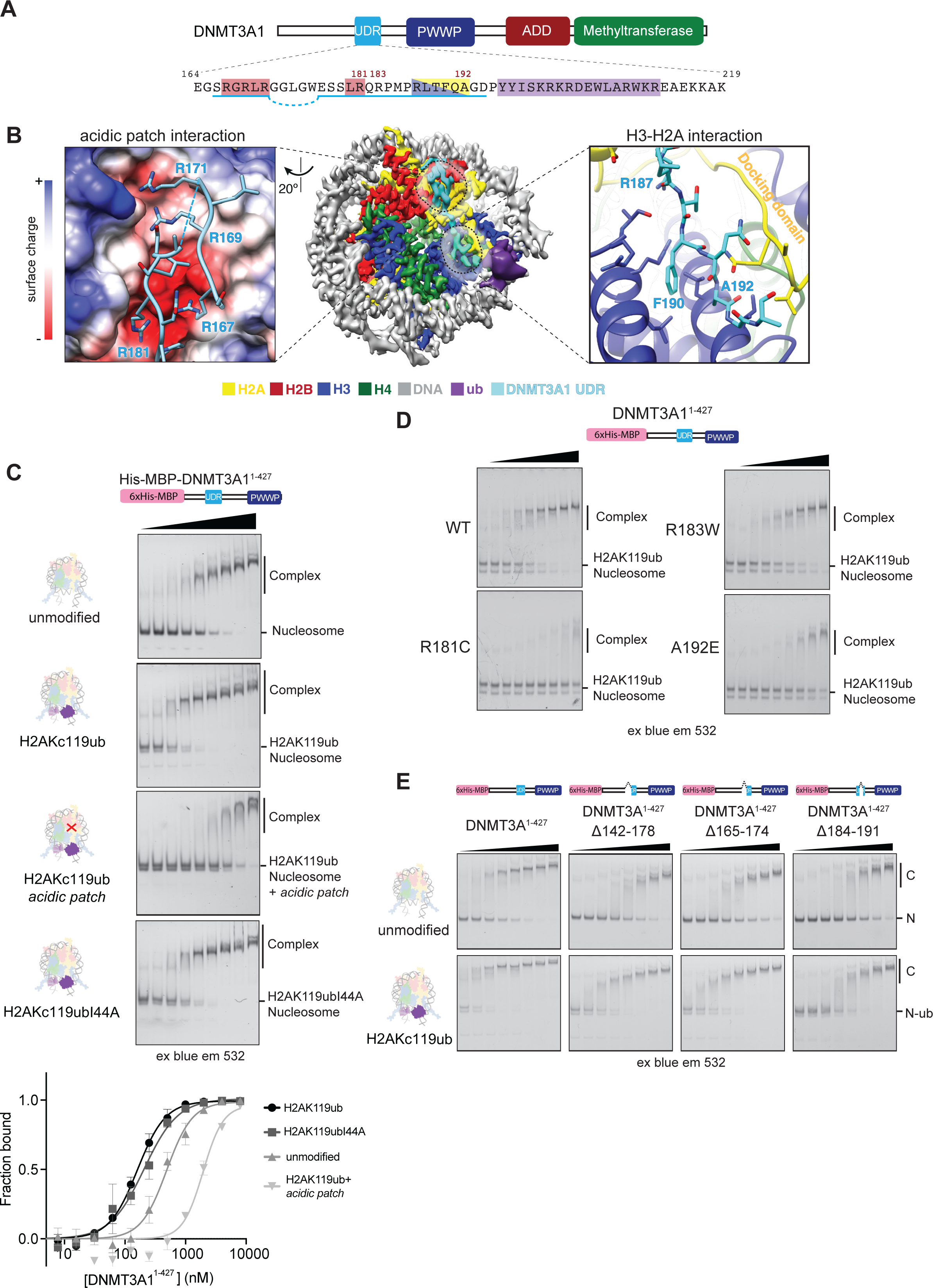
The DNMT3A1 UDR region engages with nucleosome surface. A. Schematic of DNMT3A1 and sequence of the UDR region. Ares underlined have been modelled into the density. Areas highlighted are involved in interaction with acidic patch (Moyal *et al*), H2A/H3 recess (yellow/blue) and ubiquitin (purple). B. (left) Magnified view of the H2A/H2B acidic patch-UDR interaction interface, highlighting the arginine anchor Arg181 and other positively charged residues in proximity of the acidic patch histone surface coloured according to columbic surface charge. (right) Enlarged view of H2A docking domain and H3 α2 interaction interface showing Phe190 interaction with aliphatic histone residues and H3 C-terminal interaction with backbone. C. EMSA comparing binding of DNMT3A1^1-427^ to unmodified, H2AKc119ub, H2AKc119ub acidic patch (H2A^E61A/E91A/E92A^ and H2B^E105A^) and H2AKc119ubI44A nucleosomes. Complexes were resolved by native-PAGE and imaged using blue light excitation and 532nm emission filters. Gels show concentrations 32nM-8000 nM, quantification was done with full concentration series from two experiments. D. EMSA comparing binding of DNMT3A1 cancer mutations R181C, R183W and A192E (0-8000 nM, 2x dilution series) to H2AKc119ub nucleosomes. Gels show concentrations 31nM-8000 nM for clarity experiment done in duplicate. E. EMSA comparing binding of DNMT3A1^1-427^ constructs (0-4000 nM, 1.5x dilution series) with internal deletions in the UDR to unmodified and H2AKc119ub nucleosomes. Internal deletions were created prior to (Δ142-178, Δ165-174) or after (Δ184-191) the arginine anchor Arg181 region, but still showed disruption in binding to both unmodified and ubiquitylated nucleosomes. Complexes were resolved by native-PAGE and imaged using blue light excitation and 532nm emission filters. Gels show concentrations 234nM-4000 nM for clarity from one experiment.

### The DNMT3A1 UDR recognises generic nucleosome features

In agreement with our biochemical experiments, the DNMT3A1 UDR interacts with the acidic patch region. Within the acidic patch a clear arginine anchor (McGinty & Tan, 2021) corresponding to DNMT3A1 Arg-181 projects into canonical cavity formed between H2B αC and α2/α3 helices of H2A, interacting with the carboxylate groups of Glu-61, Asp-90, and Glu-92 in H2A (Fig 5B, S7A). As seen before for interaction on H3K36me2 and unmodified nucleosomes (Fig 2C & S2C), charge reduction in the acidic patch greatly reduced binding to H2AKc119ub nucleosomes (Fig 5C & S7B). Mutation of Arg-181 and Arg-183 ablated binding to nucleosomes (Fig 2D & E) and similarly R181C greatly reduces binding to H2AKc119ub nucleosomes (Fig 5D & S7C). Extra unmodelled density suggests the UDR region loops back over the acidic patch region to further stabilise the interaction through an arginine-rich region between residues 167-171 into two adjacent negatively charged depressions behind the canonical cavity in the acidic patch. Deletions in this region reduce interaction with both unmodified and H2AKc119ub nucleosomes (Fig 5E & S7D). This acidic patch interaction is highly extensive compared to simpler arginine anchor interactions such as the LANA peptide, but not uncommon in chromatin bind proteins (McGinty & Tan, 2021).

Internal deletions in DNMT3A1^1-427^ which still maintain the Arg-181 anchor ablate both H2AKc119ub and unmodified nucleosome interaction (Fig 5E), with the most pronounced being a deletion of residues 184-191. Our structure explains this observation, as the DNMT3A1 UDR kinks down to pack tightly within a recess formed between the H2A-H2B dimer. Arg-187 projects into this recess, forming stabilising interactions with H2A Asn-89 and H3 C-terminus (Fig S7A). Interestingly this region as well as the acidic patch is divergent in nucleosomes containing the variant H2A.Z (Suto *et al*, 2000) and DNMT3A1^1-427^ binds drastically less well to H2A.Z nucleosomes (Fig S7B & E). Even ubiquitylation at residues analogous to H2A Lys 119 (H2A.Z Kc120ub) was unable to rescue robust DNMT3A1 binding, highlighting the importance of correct nucleosome interaction for ubiquitin engagement. DNMT3A1 is absent from promoter regions (Manzo *et al*., 2017), which are enriched for H2A.Z. Reduced H2A.Z promoter-adjacent nucleosome binding may help this depletion along with the repulsive effect of H3K4 methylation.

The structure shows DNMT3A1-nucleosome interactions are further stabilised by the H2A docking domain, H3 α3 helix and the C-terminal end of the α2 helix in the opposite H3’ chain (Fig 5B). Multiple predicted backbone-backbone contacts are supplemented by DNMT3A1 Phe-190 making hydrophobic contacts with H3’ Leu-109, and H3 Leu-126, and Arg-129. The C-terminal carboxylate of H3 appears to form stabilising interactions with the backbone of DNMT3A1 and Arg-187. The carbonyl group of Phe-190 and sidechain of Gln-191 are aligned to form stabilising hydrogen bonds to amines in the backbone of H2A as does the sidechain of H2A Gln-112 with DNMT3A1 Ala-192 (Fig S7A). The nucleosome interacting region of DNMT3A1 terminates with a ∼60° bend at another hydrophobic region allowed by small sidechain amino acids in Ala-192 and Gly-193, directing the UDR towards the covalently attached ubiquitin. In line with this, substitution with larger sidechain amino acids, as is found in a clinically relevant mutation A192E (Giannakis *et al*, 2016; Huang *et al*., 2022; Zehir *et al*, 2017), reduced the NT-PWWP of DNMT3A1^1-427^ interaction both with both ubiquitylated and methylated nucleosomes (Figs 5D & 2E).

### DNMT3A1 UDR-nucleosome interactions ensures specificity for H2AK119ub

The structural density attributed to ubiquitin and the rest of the DNMT3A1 UDR is weaker and spread over a larger volume than other areas of the map, preventing us from confidently building this into our model. Nevertheless, there is a degree of stabilisation due to the UDR, with more density on the nucleosome face with the UDR density compared to the non-bound ubiquitin on the anterior side of the nucleosome (Fig 6A). Accordingly, ubiquitin density appears stronger than in a H2AK119ub alone structure (Ohtomo *et al*, 2023). The remaining portion of DNMT3A1 UDR is likely also present in this region, forming interactions with the ubiquitin but not with nucleosome features.

**Figure 6.**
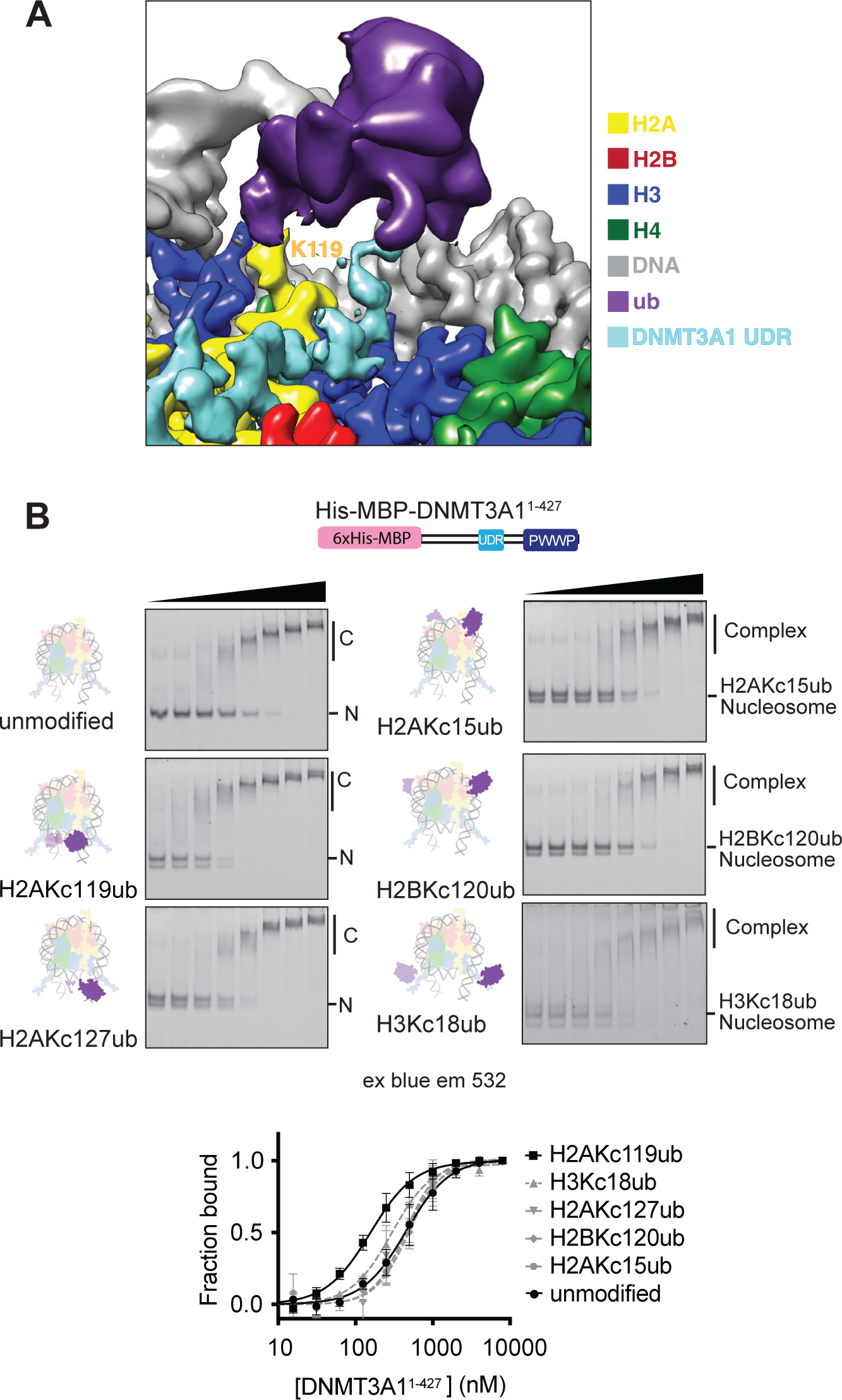
DNMT3A1 shows ubiquitin specificity and binds atypically. A. Magnified view of the H2AK119ub/UDR density from map 3, positioned above the C-terminal tail of H2A. Density from the nucleosome-contacting region of the UDR can be seen arcing upwards towards the density. Map displayed at 0.25 contour level apart from segmented ubiquitin density displayed at 0.08. B. EMSA comparing binding of DNMT3A1^1-427^ to nucleosomes ubiquitylated on different histone positions, leading to different sites on the nucleosome surface. Limiting amounts of nucleosomes were incubated with increasing concentrations (0-8000 nM, 2x dilution series) of His-MBP-DNMT3A1^1-427^. Gels show concentrations 63nM-8000 nM, quantification was done with full concentration series in triplicate.

Histone ubiquitylation is a widespread modification with different signalling outcomes based on its specific position on the nucleosome surface (Fields *et al*, 2023; Mattiroli & Penengo, 2021). DNMT3A1 co-localises with H2AK119ub-rich sites in the genome (Gu *et al*., 2022; Manzo *et al*., 2017), rather than other ubiquitylated histones. To assess the nature of ubiquitin-site specificity of DNMT3A1 we employed a chemical biology approach to make nucleosomes ubiquitylated at specific defined sites across the nucleosome surface. In addition to the facultative heterochromatic H2AKc119ub, we included DNA damage associated H2AKc15ub and H2AKc127ub (Kalb *et al*, 2014; Mattiroli *et al*, 2012), active transcription mark H2BKc120ub (Fleming *et al*, 2008; Pavri *et al*, 2006) and maintenance DNA methylation associated H3Kc18ub (Harrison *et al*, 2016; Ishiyama *et al*, 2017; Fig S8A). Ubiquitylation at other locations on the nucleosome did not influence the interaction of DNMT3A1^1-427^ in EMSA assays; the affinity of interaction was similar to that with the unmodified nucleosomes (Fig 6B). This was true even for the H2A Lys-119 proximal H2AK127ub. This suggests that despite the large size of the ubiquitin post translational modification, DNMT3A1 is a specific reader for H2AK119ub. The ubiquitin site specificity is likely mediated by the static interaction of DNMT3A1^167-193^ with the nucleosome surface, orientating and tethering the ubiquitin interacting portion of the UDR in the correct location and orientation.

The density for ubiquitin is tethered over the C-terminal tail of H2A near the linker DNA as it leaves the nucleosome core (Fig 6A). Given this proximity we tested whether interactions between linker DNA and ubiquitin may be helping to mediate DNMT3A1 interaction. If linker DNA were to be involved, we would expect higher affinity to nucleosomes containing two symmetrical linkers as both faces of the nucleosome would be bound by DNMT3A1^1-427^ rather than a single asymmetric extension. However, no ubiquitin preference was observed due to DNA length, suggesting non-nucleosomal linker DNA is not important in specifying DNMT3A1 ubiquitin specificity and interaction (Fig S8B).

### DNMT3A1 engages with ubiquitin in an atypical manner

The remaining unmodelled DNMT3A1 UDR sequence C-terminal to nucleosome interacting region is still required for H2AK119ub nucleosome interaction. We expect UDR residues 194-219 will bind to ubiquitin. In isolation the DNMT3A1 UDR region does not detectably bind to ubiquitin, neither co-eluting by size exclusion chromatography (Fig S9A) or observable in a pull-down assay (Fig S9B). Indeed, ^15^N labelled ubiquitin shows no chemical shift perturbations upon addition of high concentrations of purified unlabelled DNMT3A1 UDR sequence (Fig S9C). This suggest that the UDR region cannot be classified as a ubiquitin binding domain in isolation, such as a ubiquitin interacting motif (UIM; Komander & Rape, 2012) and can only bind within its proper nucleosomal context.

Most ubiquitin binding interactions are mediated by the canonical ubiquitin hydrophobic patch, formed by Leu-8, Ile-44, Val-70 (Komander & Rape, 2012). Surprisingly, nucleosomes harbouring H2AKc119ub with a I44A mutation could still be bound at comparable affinity by the NT-PWWP DNMT3A1^1-427^ protein (Fig 5C), suggesting the canonical ubiquitin hydrophobic patch is not implicated in H2AK119ub binding. Indeed, no clear ubiquitin interacting sequence is present in the remaining UDR region (Fig 5A, purple) overall suggesting that ubiquitin recognition by DNMT3A1 is likely atypical. Using alanine-scanning mutagenesis across the remaining UDR region shows that single mutations had no major effect on H2AKc119ub binding (Fig S9D). Taken together, the end of the UDR region interacts atypically with ubiquitin and only when it is in the correct nucleosomal context.

### H2AK119ub does not stimulate DNMT3A1 catalytic activity

Given the increased DNA methylation activity of H3K36me2 nucleosomes compared to unmodified nucleosomes observed previously (Fig 3C) we presumed that the tighter binding H2AKc119ub nucleosomes would lead to even greater enzymatic activity. Full-length DNMT3A1-DNMT3L copied the binding patterns observed for the NT-PWWP alone DNMT3A1^1-427^ fragment, with H3K36me2 and H2AK119ub nucleosomes bound preferentially over unmodified and acidic patch mutant (Fig S10A & B). Surprisingly, activity on H2AKc119ub nucleosomes was similar to unmodified nucleosomes despite significantly higher binding (Fig 7A). This disparity between affinity and catalytic activity is further exemplified when using acidic patch mutated nucleosomes which also showed equivalent activity to unmodified nucleosomes (Fig 7A). This was also observed for nucleosomes wrapped with alternative strong positioning and linker DNA sequences (Fig S10C & D) suggesting this was not an artifact of the DNA construct used. We purified DNMT3A2-DNMT3L complex (Fig S10E) to test the role of the N-terminal region in nucleosome methylation assays, absent in this isoform. DNMT3A2 also showed a similar pattern of methyltransferase activity to DNMT3A1 on the different nucleosome substrates, with higher activity on H3K36me2 nucleosome and comparable activity on H2AKc119ub nucleosome to their unmodified counterpart. This suggests that under these conditions there is a dissonance between recruitment of DNMT3A1 and downstream methylation activity (Fig 7B).

**Figure 7.**
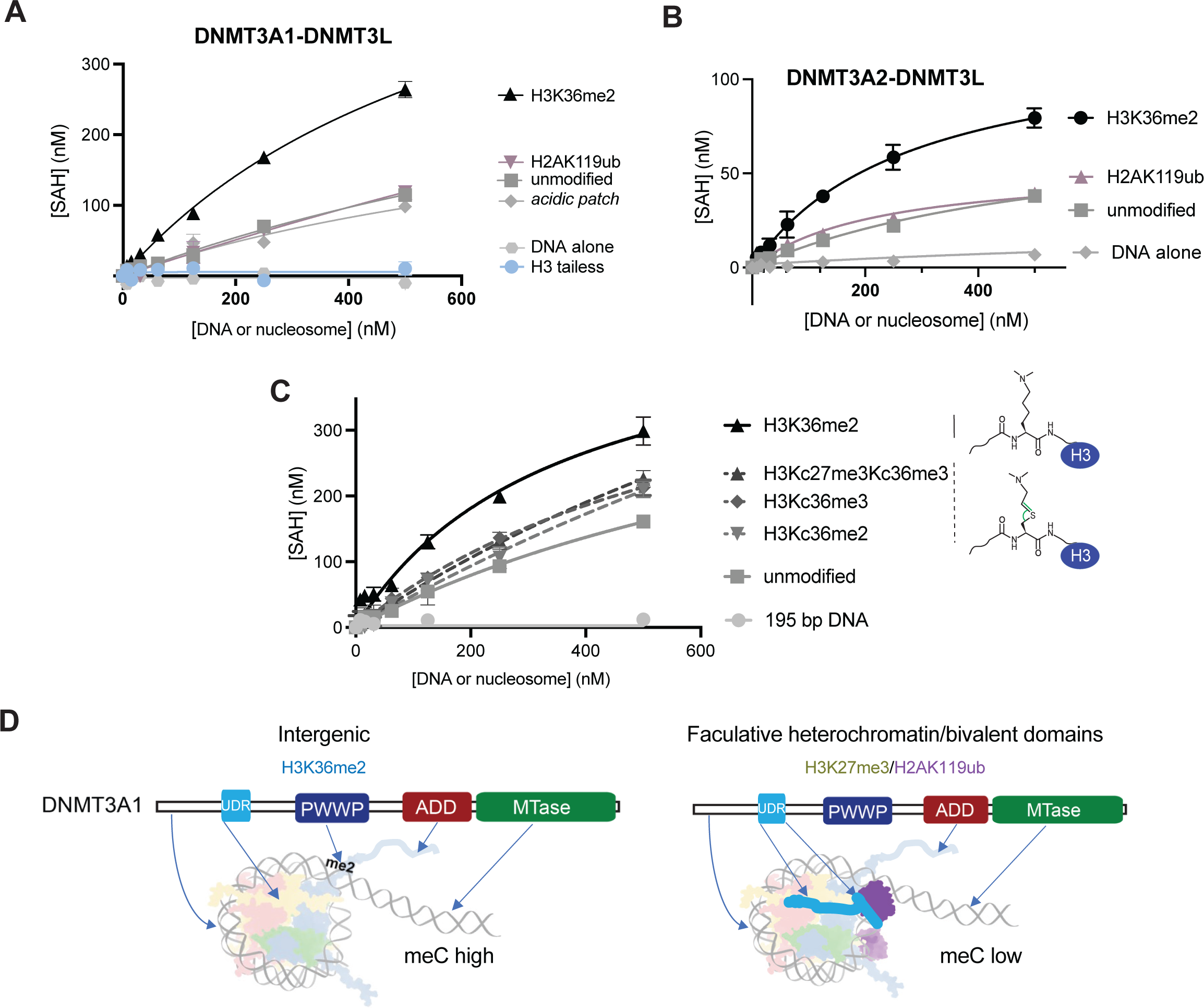
DNMT3A1 recruitment and catalytic activity are disconnected. A. Methyltransferase activity of full-length DNMT3A1-DNMT3L-StrepII on unmodified, H3K36me2, H2AKc119ub, acidic patch (H2A^E61A/E91A/E92A^ and H2B^E105A^), H3tailles (H3^A25C-136^) wrapped with 195bp Widom601 DNA and free 195bp Widom601 DNA. DNMT3A1-DNMT3L was incubated with increasing concentrations of nucleosome/DNA substrate for 1 hour at 37°C. Methyltransferase activity was detected using Promega MTase-Glo™ Methyltransferase Assay. Combined two experiments done in duplicate. Michaelis-Menten curves were fit using GraphPad Prism 10. B. Methyltransferase activity of DNMT3A2-DNMT3L-StrepII on unmodified, H3K36me2, H2AKc119ub and free 195bp DNA Widom601 DNA. Methyltransferase activity was detected using Promega MTase-Glo™ Methyltransferase Assay. Experiment done in duplicate. Michaelis-Menten curves were fit using GraphPad Prism 10. C. Methyltransferase assay comparing activity of DNMT3A1-DNMT3L on nucleosomes with different H3 methylation states. Methyltransferase activity was detected using Promega MTase-Glo™ Methyltransferase Assay. Experiment done in duplicate. Michaelis-Menten curves were fit using GraphPad Prism 10. (left) Schematic of the H3 methyl lysine analogues and methylated lysine, green highlights carbon-sulphur bond differences in bond angle and bond length compared to carbon-carbon bond. D. Proposed model summarising the results, DNMt3A1’s N-terminal extension contains a novel nucleosome interacting region that is important for conferring extra stability to chromatin interaction. This integrates with other chromatin reading domains to orchestrate correct genomic localisation. UDR region also binds to ubiquitin when placed at position 119 of H2A, increasing overall affinity and explaining how DNMT3A1 can be recruited to bivalent chromatin and facultative heterochromatin sites.

Stimulatory activity of H3K36me2 is in part due to increased availability of DNA, caused by the H3 tail-DNA interaction being disrupted (Brohm *et al*., 2022), rather than direct DNMT3A1 binding. We tested the relative importance of tail availability and recruitment in our assay, we made nucleosomes with different methylation states on the H3 N-terminal tail (Fig S10A). We already observed that H3K27me3, the DNMT3A1 non-binding methylation mark, does not stimulate catalytic activity (Fig S4B), so stimulation is not a generic feature of H3 tail methylation. Nucleosomes containing lysine methyl analogues created by cysteine alkylation (Simon *et al*, 2007) bind less well to DNMT3A1^1-427^ than closer-to-native chemically ligated nucleosomes (Fig S10F), as has been described for other methyl-lysine interactors due to differences in C-S bond length and angle (Fig 7C, right) (Seeliger *et al*, 2012). While this slight difference in chemical structure of alkylated histone affects methyl-specific reading it is unlikely to affect DNA-histone interactions. In the methyltransferase assay, this reduced direct binding to the PWWP of alkylated histones H3Kc36me2, H3Kc36me3, H3Kc27me3Kc36me3 reduces catalytic activity compared to more native H3K36me2 (Fig 7C). This suggest PWWP binding and H3 tail availability both contribute to the stimulatory effect of H3K36me2. Overall DNA methyltransferase activity is stimulated by H3K36me2 binding by the PWWP domain, but not by H2AK119ub. Indeed, DNMT3A1 is the predominant isoform present in somatic tissues while K119ub sites are commonly hypomethylated (Fu *et al*, 2020) suggesting the lower enzymatic activity may underlie the otherwise confusing observation of H2AK119ub hypomethylation.

## Discussion

Here we have shown that DNMT3A1 multivalently engages with nucleosomes, integrating multiple signals on the nucleosome including resident histone marks and nucleosome features (Fig 7D). We identified a region in the N-terminal extension of DNMT3A1 that binds to the nucleosome surface, and we propose that this plays a direct role in recruitment of DNMT3A1 at both intergenic H3K36me2 sites as well as to facultative heterochromatin/bivalent domains, amplified by co-incident recognition of the N-terminal region via ubiquitin on H2AK119.

The N-terminal region of DNMT3A1 prior to the PWWP has previously been suggested to bind generically to chromatin (Jeong *et al*, 2009; Suetake *et al*., 2011). Here we show that this is mediated by both DNA-and nucleosome-interacting regions. We further mapped the nucleosome surface interaction to DNMT3A1 residues 164-219. This coincides with a region identified by high predicted local ordering and termed the ubiquitylation-dependant recruitment motif (UDR, Weinberg *et al*., 2021). We show that this is in fact a functional UDR and the minimal region is sufficient for specific recognition of HAK119ub nucleosomes. This sequence contains a proposed canonical nuclear localisation signal (residues 202-219, Zeng *et al*., 2020) suggesting multiple functions of this region during and after DNMT3A1 is imported into the nucleus.

Intriguingly, the affinity for H2AK119ub nucleosomes appears higher than for H3K36me2 nucleosomes. This fits with the prior observation that DNMT3A1 localisation correlates with H2AK119ub-rich CpG island shores (Manzo *et al*., 2017). However, Polycomb-enriched regions are typically found to be hypomethylated (Fu *et al*., 2020). This pattern though is less clear for more differentiated cells where DNMT3A1 expression is higher (Di Croce *et al*, 2002; Mohn *et al*, 2008; Schlesinger *et al*, 2007; Weinberg *et al*., 2019). It should be noted that genomic abundance of H3K36me2 is higher but spread more evenly across the genome, as such may work to outcompete the sparser H2AK119ub mark (Weinberg *et al*., 2019). Indeed, disease mutations in the PWWP domain responsible for H3K36me2 binding leads to redistribution of DNMT3A1 (Heyn *et al*., 2019; Kibe *et al*, 2021; Sendzikaite *et al*., 2019) predominantly to H3K27me3/H2AK119ub enriched regions (Gu *et al*., 2022; Weinberg *et al*., 2021), which we propose is mediated via DNMT3A1’s UDR-H2AK119ub recognition. The action of demethylating TET enzymes potentially explain why H2AK119ub marked sites are found to be hypomethylated (Gu *et al*, 2018; Manzo *et al*., 2017; Neri *et al*, 2013), but this may also be due to a disconnect between recruitment and enzymatic activity described here. Our structure shows DNMT3A1-DNMT3L is still present on DNA but is too low resolution in this region to observe if the enzyme is in a non- or poorly-productive state. Alternatively, the high affinity of DNMT3A1-UDR may tether the enzyme to prevent processive methylation at other sites.

The DNMT3A1-DNMT3L:H2AK119ub-nucleosome structure shows that specificity for both ubiquitin and the nucleosome is contained within a short peptidic motif. Recognition of the acidic patch mediated through arginine anchors as observed for DNMT3A1 in our structure has been described for many chromatin-interacting proteins (McGinty & Tan, 2021), but the pattern of nucleosome surface contacts has not been described for other human chromatin proteins. Interestingly, the spumavirus GAG protein binds to nucleosomes (Lesbats *et al*, 2017) with a similar overall interaction interface and fold despite a different amino acid sequence (RMSD 1.25 Å), evidently an example of convergent evolution. Another acidic patch interactor is DNMT3A2-DNMT3B3 (Xu *et al*., 2020a) which deploys a different binding strategy. DNMT3B3, like DNMT3L, is a methyltransferase adaptor protein, that while non-catalytic helps to guide DNA methylation. It is a specific feature of DNMT3B3 that is used to contact the acidic patch, rigidly orientating the rest of the methyltransferase complex on adjacent DNA. In contrast, we see low ordering for the DNMT3A1-DNMT3L methyltransferase domains on adjacent DNA suggesting much greater degrees of freedom, presumably due to the flexible sequences between the UDR and rest of DNMT3A1-DNMT3L. DNMT3B3 and DNMT3A1 are almost certainly co-occurrent and likely form a tetrameric assembly (Duymich *et al*, 2016; Xu *et al*., 2020a; Zeng *et al*., 2020). How these complexes co-ordinate their nucleosome and chromatin binding features would be a fascinating avenue of future study.

Nucleosome-ubiquitin interactions mediated by short peptidic motifs or parts of domains not predicted to bind ubiquitin have been revealed in several other studies (Anderson *et al*, 2019; Fradet-Turcotte *et al*., 2013; Hsu *et al*, 2019; Rahman *et al*, 2022; Valencia-Sanchez *et al*, 2019; Wilson *et al*., 2016; Worden *et al*, 2019; Worden *et al*, 2020). Like the DNMT3A1 UDR some of these have been described to show poor to undetectable binding to ubiquitin in isolation and leverage nucleosome surface biding to ensure specific orientation of ubiquitin interacting fragments. We have previously seen that combining nucleosome tethering and ubiquitin recognition elements ensures a specific readout in other readers (Burdett *et al*., 2023; Kitevski-LeBlanc *et al*, 2017; Wilson *et al*., 2016). Indeed, while this orientation provides specificity it does not necessarily mean that the ubiquitin binding moiety is rigidly bound (Kitevski-LeBlanc *et al*., 2017; Rahman *et al*., 2022). Ubiquitin interactions are commonly mediated via the canonical hydrophobic patch (Komander & Rape, 2012), a feature that is also shared by nucleosome-ubiquitin interacting structures (Hsu *et al*., 2019; Kasinath *et al*, 2021; Wilson *et al*., 2016). Unusually, DNMT3A1’s UDR appears to interact elsewhere in ubiquitin, likely using one of the other interfaces (Komander & Rape, 2012). Further study is required to tease out the nature of this interaction.

H2AK119ub is the most prevalent histone ubiquitin mark (Fursova *et al*, 2019; Lee *et al*, 2015) and has been reported to be read by two independent ubiquitin binding domains found in the H3K27me3 methyltransferase PRC2 (Blackledge *et al*, 2014; Kasinath *et al*., 2021). Like the DNMT3A1 UDR, both of these subunits form interactions with the acidic patch. Unfortunately, the flexibility in our system prevented us from visualising the nature of the ubiquitin-DNMT3A1 interaction, but we do not see clear DNA binding of the DNMT3A1 UDR as observed for PRC2 component JARID2 nor ubiquitin displacement as observed for zinc fingers of AEBP2, suggesting these proteins have different binding modes. However, the UDR region contains a stretch of positively charged amino acids (Fig 5A), which affects recruitment to H2AK119ub when mutated (Gu *et al*., 2022). These residues may contact DNA as part of ubiquitin interaction, but based on our biochemistry this is not via linker DNA (Fig S8B). Another highly specific H2AK119ub reader SSX1 (McBride *et al*, 2020; Tong *et al*, 2023) also engages with the nucleosome acidic patch region and displays an alternate binding mode for ubiquitin recognition (Tong *et al*., 2023). As such, all 4 readers of H2AK119ub described structurally to date share common features and likely would compete for the same interfaces but display highly variable binding modes. Future work will determine the balance of affinities between these different readers and how they may integrate in a cellular setting.

Our findings have implications for the regulation of DNA methylation patterns and offer insights into the molecular basis of diseases associated with DNMT3A mutations. A previously unexplained cluster of mutations associated with developmental diseases and cancers can be found in the nucleosome contacting region of the UDR (Basturk *et al*., 2017; Dutton-Regester *et al*., 2013; Huang *et al*., 2022; Jaiswal *et al*., 2014; Tatton-Brown *et al*., 2018), with mutations affecting overall nucleosome interaction (Fig 2E, 5D). DNMT3A1 is the predominant isoform of DNA methyltransferase in the postnatal brain, which undergoes DNA methylation reconfiguration during postnatal neuronal development (Lister *et al*, 2013). Mice lacking the DNMT3A1 isoform die perinatally and this can be rescued by re-expression in the nervous system (Gu *et al*., 2022). Furthermore, aberrant localisation of DNMT3A1 affects neural development (Gu *et al*., 2022; Hamagami *et al*., 2023; Sendzikaite *et al*., 2019). Strikingly, the absence of missense mutations in the UDR region in healthy population (Fig 4B) underlie the importance of the UDR-nucleosome contacting DNMT3A1 region for normal development in humans.

## Methods

### Generation of plasmid constructs

A list of expression constructs used in this study can be found in Supp Table S2. Human DNMT3A1 and DNMT3L constructs were PCR amplified from cDNA expression constructs (Addgene #35521, #35523) and cloned by ligation dependant cloning into x6His-MBP-TEV or 6xHis-MBP-GFP expression vectors. Histone expression vectors have been described previously (Burdett *et al*., 2023; Salguero *et al*, 2019; Wilson *et al*, 2019), purchased from Addgene originated by the Landry lab. All histones are derived from human sequences, throughout all H3 constructs are based on H3.1 containing C96S C110A mutation.

Di-nucleosome DNA was designed with two strong positioning sequences (Fig S3E) and synthesised as a double-strand gBlock fragments (Integrated DNA technologies), prior to Gibson assembly into a pUC57 backbone.

DNMT3A and histone mutations were mutated either using site directed mutagenesis or direct cloning of synthesised double-strand gBlock fragments containing mutations (Integrated DNA technologies) using Gibson assembly (Gibson *et al*, 2009).

### Protein purification

#### Histone expression and purification

Histones were expressed and purified as previously described (Deak *et al*, 2023; Huntington Study *et al*, 2016; Muthurajan *et al*, 2004; Salguero *et al*., 2019). Briefly, histones were expressed in BL21 (DE3 RIL) cells and resolubilised from inclusion bodies. Histones were further purified by cation exchange chromatography prior to dialysis in 1 mM acetic acid and lyophilisation.

Histone concentrations were determined via absorbance at 280 nm using a Nanodrop One spectrophotometer (Thermo Scientific), followed by SDS-PAGE and InstantBlue (Expedeon) or Coomassie colloidal blue staining with comparison to known amounts of control proteins.

#### Expression and purification of DNMT3 constructs

His-MBP tagged DNMT3A and DNMT3B non-full-length constructs were expressed in BL21 (DE3 RIL) E. coli cells with 400 μM IPTG at 18°C overnight in 2xYT medium (16 g/l tryptone, 10 g/l yeast extract, 5 g/l NaCl, pH 7.0). Cell pellets were resuspended in in lysis buffer (25 mM sodium phosphate pH7.5, 400 mM NaCl, 0.1 % (v/v) Triton, 10% (v/v) glycerol, 2 mM β-mercaptoethanol, 1mM AEBSF, 1X protease inhibitor cocktail (2.2 mM PMSF, 2 mM benzamidine HCl, 2 μM leupeptin, 1 μg/ml pepstatin A), 4 mM MgCl2, 5 μg.mL-1 DNAse and 500 μg/ml lysozyme) and stirred at 4°C before additional lysis using a sonicator (2 s on, 2 s off for total 20 s at 50% amplitude. Cell debris was spun down at 39000 x g for 25 min and the supernatant was filtered through a 0.45 μm filter. Lysate was then loaded on Ni-NTA beads (1 ml per litre culture), washed with 25 column volume (CV) 15 mM sodium phosphate, 500 mM NaCl, 10% glycerol, 15 mM imidazole, 2 mM β-mercaptoethanol and eluted with 5CV 20 mM Tris pH7.5, 400 mM NaCl, 300 mM imidazole, 10% glycerol, 2 mM β-mercaptoethanol. The eluted protein was concentrated using a 30 kDa MWCO centrifugal filter and analysed for purity on SDS-PAGE. Where necessary, an ion exchange step was added. For this, the eluted fraction of the Nickel NTA was diluted to 100 mM NaCl using 20 mM HEPES pH7.5, 10% glycerol, 1 mM DTT and loaded on 5mL HiTrap Q HP cation exchange chromatography column (Cytiva), washed with 15 CV 20 mM HEPES pH7.5, 100 mM NaCl, 10% glycerol, 1mM DTT and eluted with 0-50% 20 mM HEPES pH7.5, 1M NaCl, 10% glycerol 1mM DTT. Fractions containing the correct protein were pooled. All proteins were further purified by size exclusion chromatography. Pooled fractions from Nickel NTA or ion exchange were concentrated to less than 5 ml and loaded on a HiLoad Superdex 200 16/600 (Cytiva) equilibrated with 15 mM HEPES pH 7.5, 150 mM NaCl, 1 mM DTT, 5% (v/v) glycerol. Fractions were analysed by SDS-PAGE and those containing pure protein were pooled, concentrated in a 30 kDa MWCO centrifugal filter, flash frozen in liquid nitrogen and stored at −80°C.

For NMR experiments His-MBP-DNMTA1^164-219^ was expressed and purified as above with additional steps to remove the tag. The His-MBP tag was cleaved using His-tagged Recombinant Tobacco Etch Virus (TEV). A ratio of 14:1 (w/w) DNMTA1^164-219^ : TEV was added to the eluted fraction of the Nickel NTA purification and dialysed in 2L 20mM HEPES pH 7.5, 150 mM, 4 mM Sodium citrate, 10% (v/v) glycerol, 1 mM DTT, 0.5 mM AEBSF using 3.5 MWCO SnakeSkin™ Dialysis Tubing at 4°C for 18 hours. The cleaved mixture was spun for 10 min at 40000 x g and supernatant was diluted to 100 mM NaCl with 20 mM HEPES pH7.5, 10% (v/v) glycerol, 1 mM DTT. The mixture was loaded on a 5mL HiTrap SP HP cation exchange chromatography column (Cytiva), washed with 10CV 20 mM HEPES pH7.5, 100 mM NaCl, 10% (v/v) glycerol, 1 mM DTT, 0.5 mM AEBSF and eluted with a gradient of 0-80% 20 mM HEPES pH7.5, 1M NaCl, 10% (v/v) glycerol, 1 mM DTT, 0.5 mM AEBSF. The fraction containing DNMTA1^164-219^ were pooled and purified further by size exclusion chromatography using a HiLoad Superdex 75 16/600 (Cytiva) in gel filtration buffer (15 mM HEPES pH 7.5, 150 mM NaCl, 1 mM DTT, 5% (v/v) glycerol). Fractions were analysed by SDS-PAGE and those containing pure protein were pooled and concentrated using a 1mL HiTrap SP FF cation exchange chromatography column (Cytiva) as follows. The pooled fractions were diluted to 100 mM NaCl using 20 mM sodium phosphate pH7.5, 5% glycerol, 1 mM DTT, washed with 5CV 20 mM sodium phosphate pH7.5, 100 mM NaCl, 5% glycerol, 1 mM DTT and eluted with 60% 20 mM sodium phosphate pH7.5, 1M NaCl, 5% glycerol, 1 mM DTT. Fractions with the highest concentration protein were pooled and dialysed into 1L 20 mM sodium phosphate pH7.5, 150mM NaCl, 5% glycerol, 1 mM DTT using 0.5 - 3 mL 3.5 MWCO Slide-A-Lyzer™ Dialysis Cassette (Thermo Fisher Scientific) at 4°C for 4 hours. The purified protein was flash frozen in liquid nitrogen and stored at −80°C.

Full-length His-MBP-DNMT3A1 or DNMT3A2 and His-GFP-DNMT3L-StrepII were co-expressed from two separate plasmids in BL21 (DE3 RIL) in 2xTY medium. Cell pellets were resuspended in lysis buffer (25 mM sodium phosphate pH7.5, 400 mM NaCl, 0.1 % (v/v) Triton, 10% (v/v) glycerol, 2 mM β-mercaptoethanol, 1mM AEBSF, 1X protease inhibitor cocktail (2.2 mM PMSF, 2 mM benzamidine HCl, 2 μM leupeptin, 1 μg.mL-1 pepstatin A), 4 mM MgCl2, 5 μg.mL-1 DNAse and 500 μg.mL-1 lysozyme) and lysed using a Constant Systems 1.1 kW TS Cell Disruptor. Cell debris was spun down at 39000 x g for 25 min and the supernatant was filtered through a 0.45 μm filter. Lysate was then loaded on a nickel sulphate charged HiTrap chelating column (Cytiva) equilibrated with 20 mM Tris pH7.5, 400 mM NaCl, 15 mM imidazole, 10%(v/v) glycerol, 2 mM β-mercaptoethanol, 0.5 mM AEBSF, washed 20 CV with the same buffer and eluted in 12 CV 20 mM Tris pH7.5, 400 mM NaCl, 400 mM imidazole, 10% (v/v) glycerol, 2 mM β-mercaptoethanol, 0.5 mM AEBSF. This was loaded on a StrepTrap HP column (Cytiva) equilibrated in streptrap buffer (20 mM HEPEs pH7.5, 150 mM NaCl, 4 mM sodium citrate, 10% (v/v) glycerol, 2 mM DTT, 0.5 mM AEBSF), washed with 15CV streptrap buffer and eluted in 2.5 mM D-desthiobiotin in streptrap buffer. The His-MBP and His-GFP tags were cleaved using TEV protease (1:25 ratio) at 4°C overnight. Cleaved proteins were loaded on a Heparin column (Cytiva) equilibrated in 20 mM HEPES pH7.5, 150 mM NaCl, 1 mM DTT, 5 % (v/v) glycerol, 0.5 mM AEBSF, washed with 15 CV buffer and eluted in 15-80% 150 mM – 1M NaCl, 20 mM HEPES pH7.5, 1 mM DTT, 5 % (v/v) glycerol, 0.5 mM AEBSF. DNMT3A1-DNMT3L-StrepII eluted at 32.5% (426 mM NaCl). The pure protein complex was concentrated using a spin concentrator and buffer exchanged to 150 mM NaCl. Purity was analysed by SDS-PAGE. An analytic amount (20 μg) was loaded on a Superdex 200 Increase 3.2/300 (Cytiva), fractions were analysed by SDS-PAGE.

#### Expression and purification of 6xHis-ubiquitinG76C

6xHis-ubiquitin G76C and 6xHis-ubiquitin I44A G76C were expressed and purified as previously described (Burdett *et al*., 2023). Briefly, ubiquitin proteins were expressed in BL21 (DE3 RIL) cells, lysed and purified using nickel sulphate charged HiTrap chelating column (Cytiva) and size exclusion chromatography (HiLoad Superdex S75 16/600 Cytiva) prior to dialysis in 1 mM acetic acid and lyophilisation.

### Histone chemical modification

#### Cysteine alkylation

Histone H3K36C, H3K27C and H3K27CK36C were alkylated as previously described (Simon *et al*., 2007; Simon & Shokat, 2012). Briefly, histones were resuspended in denaturing buffer and (2-chloroethyl)-dimethylammonium chloride reagent was added and incubated at 20°C for 2 h. The reaction was quenched with ∼650 μM β-mercaptoethanol and desalted using PD-10 columns (GE healthcare). The extent of reaction was checked using 1D intact weight ESI mass spectrometry (SIRCAMs, School of Chemistry, University of Edinburgh).

#### Native chemical ligation

Native chemical ligation was performed essentially as described (Bartke *et al*, 2010; Bryan *et al*, 2021), with some modifications. Peptides corresponding to human H3.1 residues 1-43 containing H3K36me2 and H3K27me2K36me2 with C-terminal thioesters were synthesised by Peptide Synthetics. H3 Δ1-44 T45C C110A histone was resuspended at 14 mg/ml in degassed 300mM sodium phosphate, 6M Guanidine, 100mM TCEP (Tris(2-carboxyethyl)phosphine hydrochloride) pH 7.9. Peptide was resuspended in degassed 300mM sodium phosphate, 6M Guanidine, 120mM MPAA (4-Mercaptophenylacetic acid) pH 7.9. Equal volume of the peptide and histones were mixed and incubated for 24 hours at 25°C with gentle agitation. The reaction was dialysed extensively into 7 M Urea, 25 mM Tris pH 7.5, 20 mM NaCl, 1 mM EDTA, 2mM β-mercaptoethanol and reacted products separated from unreacted histones and peptide by cation exchange chromatography eluting using a salt gradient. Reaction product was confirmed by SDS-PAGE and 1D intact weight ESI mass spectrometry (SIRCAMs, School of Chemistry, University of Edinburgh).

#### Histone ubiquitylation

Histones were chemically ubiquitylated as previously described (Burdett *et al*., 2023; Long *et al*., 2014; Wilson *et al*., 2016). Briefly, lyophilized 6xHis-ubiquitinG76C or 6xHis-ubiquitinI44AG76C and histones H2AK119C, H2AK13C, H2AK15C, H2AK127C, H2BK120C or H3K18C were resuspended, mixed, added to a solution of 1,3-dibromoacetone (DBA) in 100mM Tris pH7.5 and incubated for 1 hour on ice before quenching with 20mM β-mercaptoethanol. The ubiquitylated histones were purified by ion exchange chromatography (HiTrap SP HP column Cytiva) followed by nickel sulphate charged HiTrap chelating column (Cytiva) and dialysed in 1 mM β-mercaptoethanol, then lyophilized and stored at −20°C.

### Nucleosome formation

#### Octamer refolding

Octamers were refolded as previously described (Muthurajan *et al*., 2004; Wilson *et al*., 2016) Briefly, histones were resuspended in 20 mM Tris pH7.5, 6M guanidine, 10 mM DTT and mixed in a ratio of 1:1:1.5:1.5 H3, H4, H2A, H2B and diluted to a total concentration of 2 mg/ml. The histone mixture was dialysed to 15 mM Tris pH7.5, 2M NaCl, 5 mM β-mercaptoethanol, 1 mM EDTA. In case of His-tagged histones EDTA was omitted from this buffer. After dialysis, his-tagged octamers were purified on a 1 mL nickel sulphate charged HiTrap chelating column (Cytiva) and eluted in 15 mM Tris pH7.5, 2M NaCl, 5 mM β-mercaptoethanol, 300 mM imidazole. Then, 1 mM EDTA and 1/25 (w/w) TEV protease was added and dialysed into 15 mM Tris pH7.5, 2M NaCl, 1 mM EDTA, 5 mM β-mercaptoethanol for 18 hours to cleave the his-tag. All octamers were purified using size exclusion chromatography (HiLoad Superdex 200 16/600 or Superdex 200 Increase 10/300 GL Cytiva) in 15 mM Tris pH7.5, 2M NaCl, 1 mM EDTA, 5 mM β-mercaptoethanol. Octamer fractions were pooled, concentrated and stored in 50% (v/v) glycerol at −20.

#### PCR amplification of Nucleosome DNA

All DNA fragments for nucleosome reconstitution were generated by PCR amplification and purified as previously described (Burdett *et al*., 2023; Lowary & Widom, 1998; Wilson *et al*., 2019). A table for DNA sequences can be found in Supp Table S3. Fluorescent dyes were incorporated in the primers (IDT technologies, HPLC pure). PCR reactions using Pfu polymerase and oligonucleotides were pooled, filtered through a 0.4 μm filter, and applied to a 6 ml ResourceQ column (Cytiva) pre-equilibrated with 10 mM Tris pH 7.5 and 1 mM EDTA. The column was then washed extensively with 500 mM NaCl, before eluting across a 12 CV gradient from 500 mM NaCl to 900 mM NaCl. Fractions were analysed by native-PAGE, and fractions containing the desired product were pooled, concentrated by ethanol precipitation and resuspended in 10 mM Tris pH8.

#### Nucleosome wrapping

Nucleosomes were reconstituted as previously described (Burdett *et al*., 2023; Muthurajan *et al*., 2004), with some minor modifications. Purified octamers were incubated with DNA in 1:1.2 molar ratio and wrapped using an 18 h exponential salt reduction gradient (2M KCl to 0.2M KCl in 15 mM HEPES pH7.5, 1 mM DTT, 1 mM EDTA) and then dialysed to 15 mM HEPES pH7.5, 100 mM NaCl, 1 mM DTT, 1 mM AEBSF. Free DNA was removed from mono-nucleosomes by partial PEG precipitation, using 9% (w/v) PEG-6000 for 145bp DNA and for 175bp DNA and 9.5% PEG-6000 (w/v) and 150 mM NaCl was added. Pellets were resuspended in 15 mM HEPES pH7.5, 100 mM NaCl, 1 mM DTT, 1 mM AEBSF. The extent and purity of nucleosomes wrapping was checked by native-PAGE and SDS-PAGE analysis. Di-nucleosomes were reconstituted in the same manner as ‘mono’-nucleosomes, with the exception that a DNA:octamer ratio of 0.4:1 was used instead (Fig S3G), and no PEG precipitation was performed.

### Transmission electron microscopy sample preparation and data collection

A complex of DNMT3A1-DNMT3L-StrepII was formed with either H3Kc36me2 Di-nucleosomes wrapped with 340 bp Di-nucleosome DNA or H2AKc119ub nucleosomes wrapped with 195 bp Widom601 DNA in a 2.5:1 molar ratio protein:protomer for 1 hour on ice. Holey carbon R2/2 300 mesh grids (Quantifoil Micro Tools GmbH) were glow-discharged for 90 sec at 25 mA using a PELCO easiGlow glow discharge cleaning system. The complex was diluted with s-adenosyl methionine (SAM) in 15 mM HEPES pH7.5, 1 mM DTT to a final concentration of 110 −130 ng/μl (DNA concentration), 65 mM NaCl and 100 μM SAM and immediately vitrified by applying 3.5 μl to the glow-discharged Quantifoil grids, followed by immediate blotting (blot force = 0 N, blot time = 8 s) and plunge-freezing in liquid ethane cooled by liquid nitrogen, using a FEI Vitrobot IV (ThermoFisher) at 100% relative humidity and with a chamber temperature set at 4°C. Grids were screened for ice quality and a small dataset was collected and processed to 2D classes on a TF20 microscope (University of Edinburgh, Cryo-transmission EM facility).

#### Data collection of DNMT3A1-DNMT3L-StrepII on H2AKc119ub nucleosomes

Two separate datasets were collected on a FEI Titan Krios transmission electron microscope (ThermoFisher) operating at 300 keV equipped with a K3 camera (Gatan), using a magnification of 105 000× and a pixel size of 0.829 Å/pixel. Movies were recorded using the EPU automated acquisition software in counting super resolution mode and a total dose of 61 e−/Å^2^ over 65 frames. One dataset was 6969 movies, the second dataset was 8522 movies, defocus values ranged from −1.5 μm to −3.0 μm. Detailed information on data collection and structure refinement of DNMT3A1-DNMT3L-StrepII on H2AKc119ub nucleosomes is shown in Table S1.

#### Data collection and processing DNMT3A1-DNMT3L-StrepII on H3K36me2 di-nucleosomes

A dataset was collected on a FEI Titan Krios transmission electron microscope (ThermoFisher) operating at 300 keV, a pixel size of 0.921 Å/pixel. A total of 3636 movies were recorded using the EPU automated acquisition software on a FEI Falcon IV direct electron detector in counting mode. A dose per physical pixel/s of 4.94 was used for each exposure movie, resulting in a total electron dose of 52.69 e−/Å^2^, fractionated across 52 EPU frames and defocus values ranging from −1.2 μm to −3.0 μm. Movies were motion corrected with MotionCor2 (RELION), imported in CryoSPARC v4.31, CTF estimated using Patch CTF and 595478 particles picked. Three rounds of 2D classification yielded 193469 particles in nucleosome-like classes of which 6 classes (25706 particles) high quality Di-nucleosome classes with DNMT3A1-DNMT3L density. Sever preferred orientation prevented reliable 3D reconstruction.

### Cryo-EM Image processing

A schematic of the data processing pipeline in shown in Fig S5. All movies were motion corrected and dose weighted with MotionCor2 (RELION). For Map1 (Fig 4C & S5), data was processed in RELION 3 and 4 (Kimanius *et al*, 2021; Zivanov *et al*, 2018), except for the initial model (generated in cryoSPARC) (Punjani *et al*, 2017). The 6969 movies of the first dataset were CTF estimated using GCTF in RELION. 1 824 711 particles were picked using 2D nucleosome templates, and extracted with a box size of 384, binned by 2. Two rounds of 2D classification were performed yielding 446, 680 particles in nucleosome-like classes. 3D classification was done with 8 classes based on the ab-initio model generated from cryoSPARC processing of this dataset. One class showed high quality nucleosomes with added density (203 793 particles) and this map and particles were refined using RELION 3D auto-refine. The extra density on the linker DNA belonging to DNMT3A1-DNMT3L-StrepII clashed with the side of the box. To dela with this issue the nucleosome, rather than the whole particle, was re-cantered by re-extraction displacing all particles by 40 pixels in y direction (y −40) and a smaller box size of 320, binned by 2. An initial model was generated from these particles in RELION to check the centring of the particles and a 3D classification with three classes was done. Two classes (203 347 particles) showed good quality nucleosomes, and these were refined using RELION 3D auto-refine. A mask was created in RELION from the refined density and used for postprocessing. Postprocessing in RELION using B-factor −50, resulted in a 5.1 Å density map of DNMT3A1-DNMT3L-StrepII on a nucleosome. The local resolution across the map was estimated using RELION local resolution algorithm using a B-factor of −50 (Map 1 Fig S5). Segmentation (Fig 4C) was performed on the filtered map from the local resolution estimation using Segger v2.5.3 in UCSF Chimera (Pettersen *et al*, 2004).

For Maps 2-4 the two DNMT3A1-DNMT3L-StrepII:H2AKc119ub nucleosomes datasets were initially treated independently. Micrographs were imported into cryoSPARC (Punjani *et al*., 2017) and CTF parameters were estimated using patch CTF. ∼1,000 particles were picked manually and 2D classified to produce templates for template-based automated picking in cryoSPARC. Two rounds of 2D classification were performed to discard poorly averaged particles. The most promising classes were pooled (Dataset 1: 527, 966 particles, Dataset 2 529, 296 particles) and used for ab-initio reconstruction and separated into three (Dataset 1) or two (Dataset 2) ab-initio classes. The best class comprising 214, 344 and 330, 169 particles were merged and re-extracted with a 384-pixel voxel size. Initial ab initio job led to 391, 167 particles contributing to a high-resolution class, which yielded a map with overall 3.0 Å resolution from non-uniform refinement (Punjani *et al*, 2020), with density for ubiquitin and DNMT3A1 features, albeit weaker than core nucleosome features. To separate sample and structural heterogeneity we created a loose mask either covering the UDR region or the ubiquitin-distal DNA region. We used these as focused masks in cryoSPARC 3D classification, with 2 classes and 0.85 similarity score (UDR density) or 4 classes and 0.5 similarity score (Ub-DNA density). For the UDR based mask on map corresponding to 191, 397 particles showed the clearest density for this region and homogenous refinement using a wide dynamic mask of 10-18 Å, produced Map 2 showing density for UDR, ubiquitin, extra-nucleosomal DNA and some poor extra density expected to be from DNMT3A1-DNMT3L. With the same particles non-uniform refinement with local CTF optimisation produced a 3.1 Å Map 3. Focused classification of the Ubiquitin density revealed poorer overall density and no discrete ubiquitin states. Pooling 2 classes followed by non-uniform refinement produced map 4. Local resolution was estimated in cryoSPARC and final maps filtered using a B factor of −50.

### Model building

An initial model of DNMT3A1 and nucleosome was generated using ModelAngelo (Jamali *et al*, 2023), using the protein sequence for human histones, 195bp DNA and residues 164-219 of DNMT3A1 with final 3.1 Å map 3 sharpened using a B factor of −50. This reliably built density into the unassigned UDR region with register that fit with biochemical observations. For model building the cryo-EM structure of human nucleosomes (PDB ID: 7XD1) (Ai *et al*., 2022) was combined with the DNMT3A1 UDR model and rigid-body docked into the reconstructed density in Chimera (Pettersen *et al*., 2004). DNA ends were removed on one side of the nucleosome, to represent the available density. The model was adjusted using Coot (Emsley *et al*, 2010) and ISOLDE (Croll, 2018) and extra residues added to unmodelled density. The model was iteratively refined using Phenix real space refine (Liebschner *et al*, 2019). Where density was lacking side chains were removed past the Cβ position. The overall quality of the model was assessed using MolProbity (Williams *et al*, 2018) and Phenix validation tools. All figures were prepared in Chimera or ChimeraX (Pettersen *et al*., 2004; Pettersen *et al*, 2021)

### Electrophoretic mobility shift assay

Nucleosomes wrapped with 6-carboxyfluorescein (5′ 6-FAM) labelled 175bp Widom601 DNA or free 6-carboxyfluorescein (5′ 6-FAM) labelled DNA were incubated at a concentration of 2.4 nM with various concentration ranges of proteins, as specified in figure legends, in EMSA buffer 15 mM HEPES pH7.5, 75 mM NaCl, 0.05% (v/v) TritonX100, 0.05 mg/mL BSA, 10% (v/v) glycerol, 1 mM DTT, 8% (w/v) sucrose, 0.01% (w/v) bromophenol blue) to a final volume of 12 μl. Competitor DNA was used in EMSAs with nucleosomes (0.5 mg/mL salmon sperm DNA, low molecular weight 31149-10g-f, Sigma-Aldrich). Samples were incubated on ice for 1 h and products were separated on 5% 19:1 acrylamide native PAGE gels using 1xTris Glycine as running buffer for 90 min at 4°C. Gels were imaged for FAM signal (Excitation Blue light, Emission 532nm) using Bio-Rad ChemiDoc MP. Quantification was done using Image Lab (Bio-Rad) and binding curves were analysed in GraphPad Prism 10 using non-linear regression – specific binding with hill slope.

For full-length protein nucleosomes wrapped with 195 bp Widom601 DNA or free 195 bp Widom601 DNA as used in methyltransferase assays were incubated at a concentration of 2.4 nM with a concentration range of 0-2.6 µM, 2x dilution series, in 17.5 mM HEPES pH8, 150 mM NaCl, 2.5% (v/v) glycerol, 1 mM DTT, 0.1 mg/mL BSA, 1.5 mM MgCl2, 8% (w/v) sucrose, 0.01% (w/v) bromophenol blue. Samples were incubated on ice for 1 h and products were separated on 5% 19:1 acrylamide native PAGE gels using 1xTris Glycine as running buffer for 90 min at 4°C. Gels were stained with Diamond stain (Promega) and imaged on a Bio-Rad ChemiDoc MP.

### Nuclear Magnetic resonance (NMR)

^1^H-^15^N HSQC NMR of ^15^N-labelled ubiquitin was performed with and without DMT3A^164-219^ in identical normalised buffer (20 mM sodium phosphate pH 7.5, 150 mM NaCl, 5% (v/v) glycerol, 1 mM DTT). Spectra were taken of ^15^N ubiquitin (300 μM) and then DNMT3A1^164-219^ was titrated in to a final concentration of 300 μM DNMT3A1^164-219^ and 125 μM of ubiquitin, at a molar ratio of 2.4 : 1. Titrations of DNMT3A1^164-219^ were also tested to monitor any dose response shifts. All spectra were measured using a Bruker Avance NEO 800 MHz standard bore NMR spectrometer with Topspin (Bruker) software, at 298K, with 32 scans, sweep width 12 ppm in the ^1^H dimension and 35 ppm in ^15^N dimension. Spectra were analysed using CCPN mr3.1.1 AnalysisAssign and the crosspeaks were assigned based on BMRB17769 (Cornilescu *et al*, 1998) and previous ubiquitin spectra.

### Nucleosome Pull-down assay

Amylose resin (NEB) was equilibrated with amylose pull down buffer (50 mM Tris pH7.5, 150 mM NaCl, 0.02% (v/v) NP-40, 10% (v/v) glycerol, 1 mM EDTA, 0.1 mg/mL BSA and 2 mM β-mercaptoethanol) and blocked with 1 mg/mL BSA in amylose pull down buffer. After washing 3x with amylose pull down buffer, His-MBP tagged proteins (6 µg unless differently specified) were immobilised on amylose beads for 1 hour rotating at 4°C and washed 3x with amylose pull down buffer. Nucleosomes (0.5 μg unless differently specified) were added and incubated for 2 hours at 4°C, washed with amylose pull down and resuspended in 30 μl 2x SDS loading buffer. Samples (6 μl) and 5% input were loaded on SDS-PAGE gels. Nucleosomes bound were detected using western blot followed by detection with histone antibodies (anti-H3 Abcam ab1791; anti-H2A Abcam ab18255, anti-H2B Abcam ab1790) and HRP conjugated secondary antibodies (anti-mouse IgG-HRP Vector Labs PI2000, anti-rabbit IgG-HRP Vector Labs PI1000). Proteins were detected using anti-MBP antibody (anti-MBP NEB e8032s) and HRP conjugated secondary antibody, PonceauS (3% trichloroacetic acid, 3% sulfosalicylic Acid, 0.2% Ponceau) or stainfree gels (Mini-PROTEAN TGX Stain-Free Precast Gels).

For the alanine scanning experiment partially purified His-MBP-DNMT3A1^1-427^ and variants were expressed in small scale and enriched from *E. coli* extract with a Ni-NTA bead purification step, as described above under protein purification. Protein was eluted and concentration determined, prior to loading on amylose beads at ∼ 5 μg per pulldown.

### DNA methyltransferase assays

Methyltransferase activity of full-length DNMT3A1-DNMT3L and DNMT3A2-DNMT3L was measured using Promega MTase-Glo™ Methyltransferase Assay following the small volume (10 μl) protocol with minor adjustments. Nucleosomes wrapped with 195bp Widom 601 DNA (0-0.5 µM) and S-adenosyl methionine (SAM, 20 μM) were incubated with full-length DNMT3A1-DNMT3L (0.5 μM) in 20 mM HEPES pH8, 150 mM NaCl, 3 mM MgCl2, 0.1 mg/ml BSA, 1 mM DTT, 0.25% glycerol for 1 hour at 37°C. 5x reaction mixture (Promega MTase-Glo™ Methyltransferase Assay) was added and the mixture was incubated for 30 min at room temperature followed by 30 min incubation with detection reagent (Promega MTase-Glo™ Methyltransferase Assay). Luminescence was detected using a SpectraMax iD5 (Molecular Devices) plate reader.

A SAH standard curve was made (0-1μM SAH) in 20 mM HEPES pH 8, 150 mM NaCl, 3 mM MgCl2, 0.1 mg/ml BSA, 1 mM DTT, 0.25% (v/v) glycerol following the same steps as the methyltransferase reaction. Raw data were baseline subtracted and converted using the slope of the SAH standard curve. Michaelis-Menten curves were fit using Graphpad Prism 10.

### Missense variants analysis

Missense variant analysis was performed as described before (Deak & Cook, 2022). Briefly, data on missense variants associated with the human DNMT3A1 gene (transcript ENST00000264709.3, genome build GRCh37 / hg19) were retrieved from the gnomAD v2.1.1 dataset (Karczewski *et al*., 2020). Plot Protein Converter (Deak & Cook, 2022) was used to filter the data for non-deleterious variants and format it for the Plot Protein program (Turner, 2013), enabling visualisation of variants on the DNMT3A1 protein sequence.

### Ubiquitin pull down assays

Amylose resin (NEB) was equilibrated with amylose pull down buffer (50 mM Tris pH7.5, 150 mM NaCl, 0.02% (v/v) NP-40, 10% (v/v) glycerol, 1 mM EDTA, 0.1 mg/mL BSA and 2 mM β-mercaptoethanol) and blocked with 1 mg/ml BSA in amylose pull down buffer. After washing 3x with amylose pull down buffer, His-MBP and His-MBP-DNMT3A^1-427^ (50 µg) were immobilised on amylose beads for 1 hour rotating at 4°C and washed 3x with amylose pull down buffer. His-TEV-UbiquitinG76C (50 μg) was added and incubated for 2 hours at 4°C, washed with amylose pull down and resuspended in 30 μl 2x SDS loading buffer. Inputs of ubiquitin, His-MBP and His-MBP-DNMT3A1^1-427^ (2%) were loaded as control. Proteins were detected using stainfree gels (Mini-PROTEAN TGX Stain-Free Precast Gels), ubiquitin was detected using anti-ubiquitin antibody (Santa Cruz sc-8017) and HRP conjugated secondary antibody.

### Analytical Ubiquitin and DNMT3A^1-277^ coelution assay

DNMT3A1^1-277^ (20 μg) with or without ubiquitin in a molar ratio of 1:5 was incubated on ice for 1 hour at a concentration of 4 mg/ml. The protein or protein complex was diluted in size exclusion buffer to 2 mg/mL and loaded on a Superdex increase S200 3.2/300 column equilibrated with 20 mM HEPES pH 7.5, 150 mM NaCl, 1 mM DTT, 5% (v/v) glycerol. Eluted peak fractions were analysed on SDS-PAGE gels.

### Crosslinking Mass spectrometry

A complex of DNMT3A1^1-427^ and H3Kc36me2 modified nucleosomes (2.5:1 molar ratio) was crosslinked using 1-ethyl-3-(3-dimethylaminopropyl)carbodiimide hydrochloride (EDC) and N-hydroxysulfosuccinimide in a w / w ratio of 1 : 15-30 : 30-60 nucleosome : EDC : sulfo-NHS in 50 mM HEPES pH7.5, 150 mM NaCl, 1 mM DTT. The complex was crosslinked for 4 hours on ice then quenched with 50 mM Tris pH7.5. Crosslinked proteins were separated on an SDS PAGE gel. The bands running at a higher molecular weight than DNMT3A1^1-427^ were excised and the proteins were digested following previously established protocol (Maiolica *et al*, 2007). Briefly, proteins were reduced with 10 mM DTT for 30min at 37°C, alkylated with 55 mM iodoacetamide for 20min at room temperature and digested using 13 ng/μl trypsin (Promega) overnight at 37°C. Digested peptide were desalted using C18-StageTips (Rappsilber *et al*, 2003; Rappsilber *et al*, 2007) for LC-MS/MS analysis.

LC-MS/MS analysis was performed using Orbitrap Fusion Lumos (Thermo Fisher Scientific) with a “high/high” acquisition strategy. The peptide separation was carried out on an EASY-Spray column (50 cm × 75 μm i.d., PepMap C18, 2 μm particles, 100 Å pore size, Thermo Fisher Scientific). Mobile phase A consisted of water and 0.1% v/v formic acid. Mobile phase B consisted of 80% v/v acetonitrile and 0.1% v/v formic acid. Peptides were loaded at a flow rate of 0.3 μl/min and eluted at 0.2 μl/min using a linear gradient going from 2% mobile phase B to 40% mobile phase B over 139 min (each sample has been running three time with different gradient), followed by a linear increase from 40% to 95% mobile phase B in 11 min. The eluted peptides were directly introduced into the mass spectrometer. MS data were acquired in the data-dependent mode with 3 seconds acquisition cycle. Precursor spectrum was recorded in the Orbitrap with a resolution of 120,000. The ions with a precursor charge state between 3+ and 8+ were isolated with a window size of 1.6 m/z and fragmented using high-energy collision dissociation (HCD) with collision energy 30. The fragmentation spectra were recorded in the Orbitrap with a resolution of 30,000. Dynamic exclusion was enabled with single repeat count and 60 seconds exclusion duration.

The mass spectrometric raw files were processed into peak lists using ProteoWizard (version 3.0) (Kessner *et al*, 2008), and cross-linked peptides were matched to spectra using Xi software (version 1.7.6.1) (Mendes *et al*, 2019; Xi search) with in-search assignment of monoisotopic peaks (Lenz *et al*, 2018). Search parameters were MS accuracy, 3 ppm; MS/MS accuracy, 10ppm; enzyme, trypsin; cross-linker, EDC; max missed cleavages, 4; missing mono-isotopic peaks, 2; fixed modification, carbamidomethylation on cysteine; variable modifications, oxidation on methionine and phosphorylation on threonine for phosphorylated sample; fragments, b and y ions with loss of H2O, NH3 and CH3SOH. K36Cme2 at H3.1 sequence using Customer setting modification to define.

### Data Availability

The cryo-EM density map, and associated meta data for the NCP deposited at the Electron Microscopy Data Bank under accession number EMD-18778 & EMD-XXX, and structural model under PDB 8QZM. Raw micrograph data is available at EMPIAR-XXX.

Crosslinking mass spectrometry data has been uploaded to PRIDE database under XXX.

## Acknowledgements

We thank Duncan Sproul, Adrian Bird and members of the Wilson lab for helpful discussions and critical reading of the manuscript. MDW’s work is supported by the Sir Henry Dale Fellowship form the Wellcome Trust [210493/Z/18/Z], Medical Research Council (T029471/1), Scottish high-field NMR centre seed funding and University of Edinburgh. HW work is part supported by Institutional Strategic Support Fund (IS3-R1.37 22-23). Work in the Voigt lab was supported by the Wellcome Trust ([104175/Z/14/Z], Sir Henry Dale Fellowship to P.V.) and the UK Biotechnology and Biological Sciences Research Council (BBS/E/B/000C0421). D.V, A.D, & D.K are supported by PhD studentships from the Darwin Trust of Edinburgh. W.R and J.A.W are funded by the Wellcome trust integrative cellular mechanisms PhD program (218470). This work was supported by the Edinburgh Protein Production Facility (EPPF), which receives funding from a core grant (203149) to the Wellcome Centre for Cell Biology at the University of Edinburgh. Grid screening was performed in the cryo-EM facility in School of Biological Sciences at the University of Edinburgh, we are grateful to Maarten Tuijtel and Martin Singleton for their support. The cryo-EM facility was set up with funding from the Wellcome Trust (087658/Z/08/Z) and SULSA. We are grateful to the Rappsilber lab for access to the Xi server for mapping protein crosslinks. We thank Arthur Riggs, Joe Landry (via Addgene) Duncan Sproul and Frank Sicheri for gifts of plasmids. We are very grateful to David Owen, Julika Radecke and Matt Byrne at Diamond for access and support of the Cryo-EM facilities at the UK national electron bio-imaging centre (eBIC), proposal EM-BI24557 & EM-BI31827, funded by the Wellcome Trust, MRC and BBSRC. This work was supported by the Wellcome Centre for Cell Biology Mass spectrometry facility which receives funding from a core grant to the Wellcome Centre for Cell Biology (203149) and a Multi-User Equipment grant(s) (108504). We thank Logan Mackay in SIRCAMS school of chemistry, university of Edinburgh for mass spec analysis. We thank Juraj Bella in the NMR facility, School of Chemistry, University of Edinburgh for help with NMR experiments.

## Author contributions

MDW conceived the study and supervised the project. HW and MDW designed the experiments (unless otherwise stated), analyzed the data, and wrote the manuscript, with input from the other authors. MRDT, GD, YZ, DK, JAW, JM, AD, MDW, DV, PV and HW purified protein and DNA components. Sample loading and data analysis of Crosslinking mass spectrometry was performed by JZ. Cryo-EM image processing was performed by MDW and HW. Model building was performed by MDW and HB. Biochemical assays were performed by MDW, GC, WR, and HW. NMR analysis was done by HW and JB.

---

**Supplemental Table 1.**
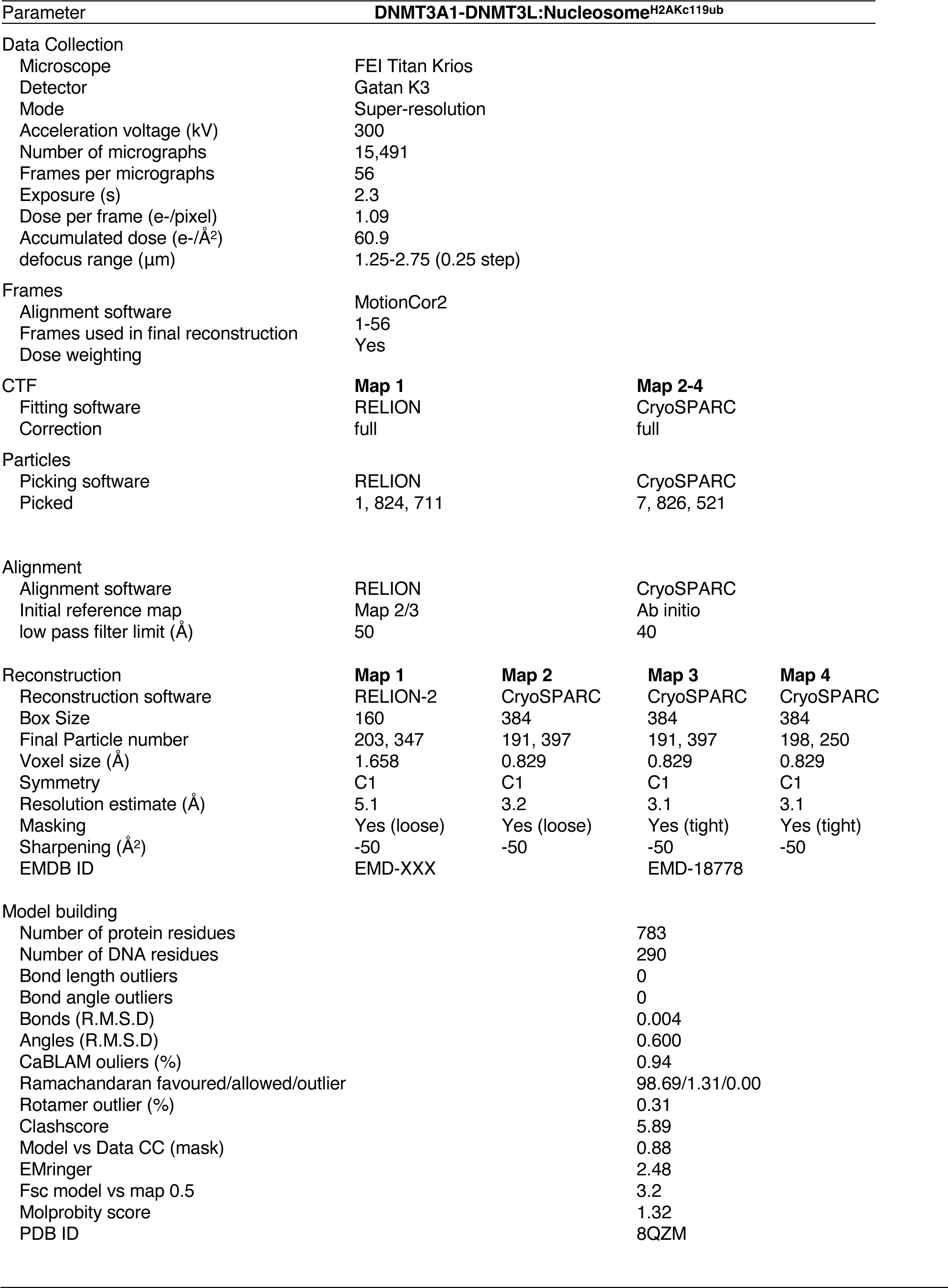
Cryo-EM data collection, refinement and validation statistics.

**Supplemental Table 2.** Expression constructs used in this study.

| Construct | Protein (aa) | Domain/regions(s) | Mutations |
| --- | --- | --- | --- |
| His-MBP |  |  | - |
| His-MBP-DNMT3A1-427 | 1-427 | NT UDR PWWP | - |
| His-MBP-DNMT3A1-277 | 1-277 | NT UDR | - |
| His-MBP3A278-427 | 278-427 | PWWP | - |
| His-MBP-DNMT3A1-614 | 1-614 | NT UDR PWWP<br>ADD | - |
| His-MBP-DNMT3A1-482 | 1-482 | NT UDR PWWP | - |
| His-MBP-DNMT3A1-427 K53A/K55A | 1-427 | NT UDR PWWP | K53A/K55A |
| His-MBP-DNMT3A1-427 R181A/R183A | 1-427 | NT UDR PWWP | R181A/R183A |
| His-MBP-DNMT3A1-427 K53/K55/R181/R183A | 1-427 | NT UDR PWWP | K53A/K55A/<br>R181A/R183A |
| His-MBP-DNMT3A1-427 R181C | 1-427 | NT UDR PWWP | R181C |
| His-MBP-DNMT3A1-427 R183W | 1-427 | NT UDR PWWP | R183W |
| His-MBP-DNMT3A1-427 A192E | 1-427 | NT UDR PWWP | A192E |
| His-MBP-DNMT3A1-427 W330R | 1-427 | NT UDR PWWP | W330R |
| His-MBP-DNMT3A1-427 D333N | 1-427 | NT UDR PWWP | D333N |
| His-MBP-DNMT3A1-427 P195A | 1-427 | NT UDR PWWP | P195A |
| His-MBP-DNMT3A1-427 Y197A | 1-427 | NT UDR PWWP | Y197A |
| His-MBP-DNMT3A1-427 I198A | 1-427 | NT UDR PWWP | I198A |
| His-MBP-DNMT3A1-427 K200A | 1-427 | NT UDR PWWP | K200A |
| His-MBP-DNMT3A1-427 R201A | 1-427 | NT UDR PWWP | R201A |
| His-MBP-DNMT3A1-427 K202A | 1-427 | NT UDR PWWP | K202A |
| His-MBP-DNMT3A1-427 R203A | 1-427 | NT UDR PWWP | R203A |
| His-MBP-DNMT3A1-427 D204A | 1-427 | NT UDR PWWP | D204A |
| His-MBP-DNMT3A1-427 E205A | 1-427 | NT UDR PWWP | E205A |
| His-MBP-DNMT3A1-427 L207A | 1-427 | NT UDR PWWP | L207A |
| His-MBP-DNMT3A58-427 | 58-427 | NT UDR PWWP | - |
| His-MBP-DNMT3A115-427 | 115-427 | NT UDR PWWP | - |
| His-MBP-DNMT3A142-427 | 142-427 | NT UDR PWWP | - |
| His-MBP-DNMT3A192-427 | 192-427 | NT PWWP | - |
| His-MBP-DNMTA142-219 | 142-219 | UDR | - |
| His-MBP-DNMTA164-219 | 164-219 | UDR | - |
| His-MBP-DNMT3A1-427 Δ142-178 | 1-427 | NT PWWP | Δ142-178 |
| His-MBP-DNMT3A1-427 Δ165-174 | 1-427 | NT PWWP | Δ142-174 |
| His-MBP-DNMT3A1-427 Δ184-191 | 1-427 | NT PWWP | Δ142-191 |
| His-MBP-DNMT3A1 | 1-912 | NT PWWP ADD<br>Mtase | - |
| **DNMT3A2** |  |  |  |
| His-MBP-DNMT3A21-238 | 1-238 | PWWP | - |
| His-MBP-DNMT3A2 | 1-689 | PWWP ADD Mtase | - |
| **DNMT3B** |  |  |  |
| His-MBP-DNMT3B1 1-350 | 1-350 | NT PWWP | - |
| **DNMT3L** |  |  |  |
| His-GFP-DNMT3L-StrepII | 1-386 | ADD Mtase-like | - |
| **Histones** |  |  |  |
| unmodified H3.1 (No cys) | 1-136 |  | C110A, C96S |
| H3.3 T45CΔ1-44 | 44-136 |  | T45CΔ1-44, C110A |
| H3.1 K36C | 1-136 |  | C110A, C96S, K36C |
| H3.1 K27C | 1-136 |  | C110A, C96S, K27C |
| H2A | 1-130 |  | - |
| H2A acidic patch | 1-130 |  | E61A E91A E92A |
| H2B | 1-126 |  | - |
| H2B acidic patch | 1-126 |  | E105A |
| H4 | 1-103 |  | - |
| H2A K13C | 1-130 |  | K13C |
| H2A K15C | 1-130 |  | K15C |
| H2A K119C | 1-130 |  | K119C |
| H1A K127C | 1-130 |  | K127C |
| H2B K120C | 1-126 |  | K120C |
| H3 K18C | 1-136 |  | C110A, C96S, K18C |
| H2AK119C acidic patch | 1-130 |  | E61A E91A E92A<br>K119C |
| H3tailless | 25-136 |  | C110A, C96S, A25C<br>Δ1-24 |
| H3K27CK36C | 1-136 |  | C110A, C96S,<br>K27C/K36C |
| **Ubiquitin** |  |  |  |
| His-TEV-Ubiquitin | 1-76 |  | G76C |
| His-TEV-Ubiquitin I44A | 1-76 |  | I44A/G76C |

**Supplemental Table 3.**
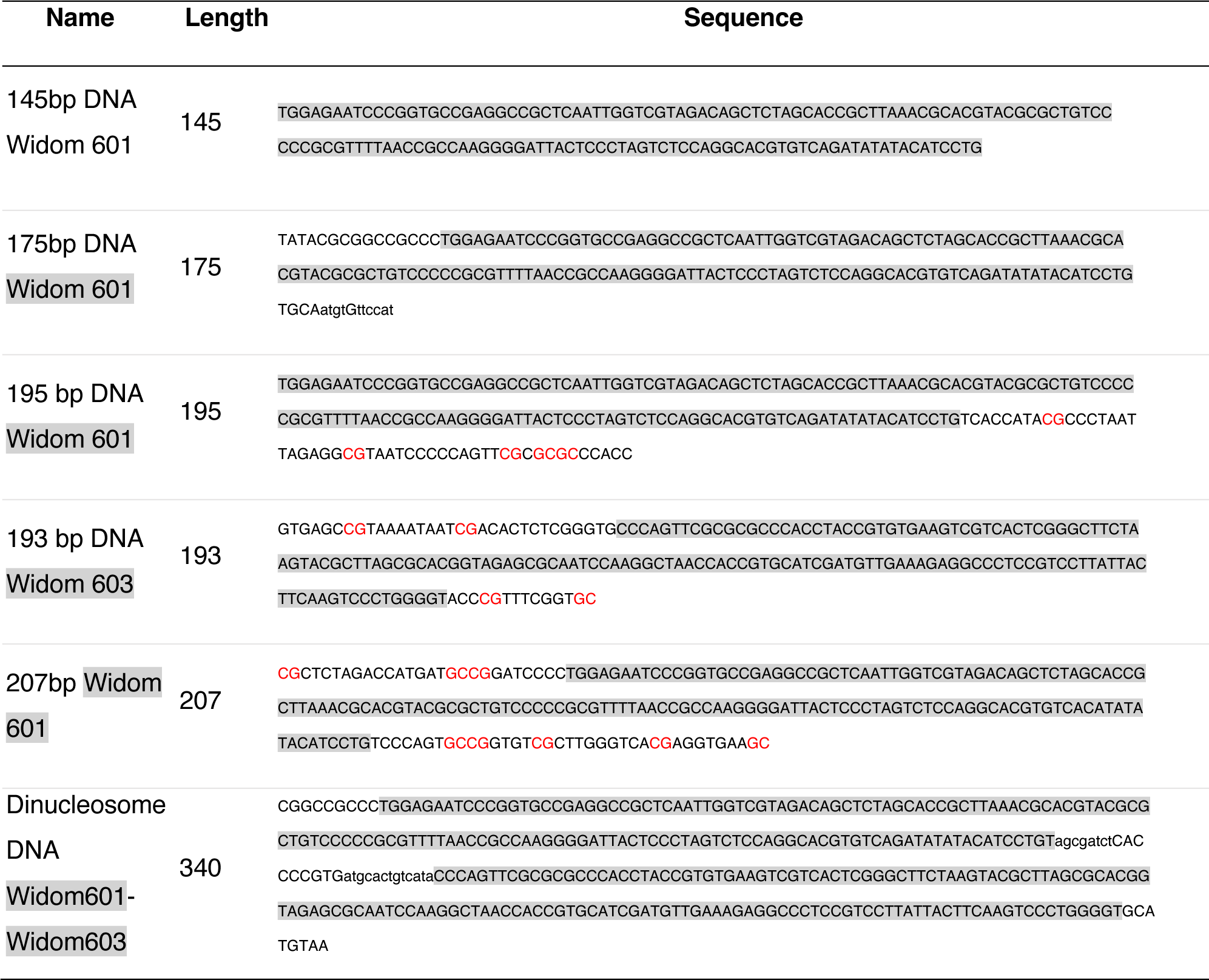
DNA sequences used for wrapping nucleosomes.

## Supplemental Figure legends

**Supplemental Figure S1. DNMT3A1-H3K36me2 nucleosome interaction is localised in N-terminus and PWWP domains.**

A. SDS-PAGE gels showing purified protein constructs used in Fig 1 and Fig 2.

B. Native gels and SDS-PAGE gels of nucleosomes used in Fig 1. Nucleosomes (100ng based on DNA concentration were loaded on 5% 19:1 acrylamide native gels using native loading buffer (8% sucrose, 0.02 mg/mL BSA, 0.01% bromophenol blue) and run 100V for 90 min. For SDS-PAGE, 900 ng nucleosomes were loaded on 17% 37.5:1 acrylamide gels using SDS-PAGE loading buffer (56 mM Tris pH6.8, 11% glycerol, 2.4%SDS, 0.016 % bromophenol blue) and run 100V for 10 min then 200V for approximately 60 min.

C. Quantification of EMSA in Fig 1B.

D. EMSA comparing DNMT3A1 ^1-427^ and DNMT3A2 ^1-238^. Limiting amounts (2.3 nM) of H3K36me2 nucleosomes were incubated with increasing concentrations (0-8000 nM) of protein. Complexes were resolved by native-PAGE and imaged using blue light excitation and 532nm emission filters. Gels show concentrations 32nM-8000 nM.

E. EMSA of DNMT3A1^1-427^ WT, W330R and D333N binding (0 - 8000 nM 2x dilution series) to unmodified and H3K36me2 nucleosomes. Gels show concentrations 18.75 - 2000 nM for clarity, quantification was done with full concentration series wild type in duplicate, and mutants a single experiment.

**Supplemental Figure S2. Mapping DNMT3A1 N-terminal interactions to the acidic patch.**

A. SDS-PAGE gel of crosslinked complex for crosslinking mass spectrometry in Fig 2A. Everything above non-crosslinked DNMT3A1 was excised from the gel and trypsin digested prior to identification in mass spectrometry.

B. Crosslinking mass spectrometry replicates from Fig 2A. Circular representation of two biological replicates one measured in triplicate (repeat 1, 2, 3). Only crosslinks between DNMT3A1 and histones are shown.

C. EMSA comparing binding of DNMT3A1^1-427^ to unmodified or acidic patch mutated (H2A^E61A/E91A/E92A^ and H2B^E105A^) nucleosomes wrapped with 5’ FAM labelled 175bp Widom601 DNA (FAM = fluorescein). Limiting amounts (2.3 nM) of H3K36me2 nucleosomes were incubated with increasing concentrations (0-4000 nM) of His-MBP-DNMT3A1^1-427^. Complexes were resolved by native-PAGE and imaged using blue light excitation and 532nm emission filters. Gels show concentrations 500nM-4000 nM for clarity.

D. EMSA comparing binding of DNMT3A1 constructs with different lengths of the N-terminal region to H3K36me2 nucleosomes wrapped with 5’ FAM labelled 175bp Widom601 DNA (FAM = fluorescein). Limiting amounts (2.3 nM) of H3K36me2 nucleosomes were incubated with increasing concentrations (0-8000 nM) of His-MBP-DNMT3A1 constructs. Complexes were resolved by native-PAGE and imaged using blue light excitation and 532nm emission filters. Gels show concentrations 32nM-8000 nM for clarity, quantification was done with full concentration series in duplicate.

E. SDS-PAGE gel of purified DNMT3A proteins used in Fig 2.

F. Pull down experiment comparing binding of DNMT3A1 with mutations in the N-terminus to H3Kc36me2 nucleosomes wrapped with 175bp Widom601 DNA. Proteins were immobilised on amylose beads and nucleosomes were added. Bound nucleosomes were detected using western blot followed by detection with antibodies against histones H3. Bound proteins were detected using an anti-MBP antibody.

**Supplemental Figure S3. Preparation of full-length DNMT3A1-3L and nucleosomes for enzymology and Cryo-EM.**

A. Schematic of the purification strategy of full-length DNMT3A1-DNMT3L-StrepII complex used. Full-length His-MBP-DNMT3A1 and His-GFP-DNMT3L-StrepII were co-expressed in E. coli BL21 RIL. Cells were lysed and purified with Nickel NTA affinity purification followed by StrepTrap. His-MBP and His-GFP tags were cleaved using TEV protease followed by a further purification using a heparin column. The salt was reduced to 150 mM NaCl and complex was flash frozen and stored at −80.

B. A sample of 40 μg of DNMT3A1-DNMT3L-StrepII (1.77mg/mL) complex was run on analytical size exclusion chromatography to assess monodispersity. Two overlapping peaks (0.9ml and 1.05ml elution volume) corresponding to rough sizes for heterotetramer and heterodimeric species. An SDS-PAGE gels was run of all fractions across the peaks.

C. Schematic of Promega MTase-Glo™ Methyltransferase Assay

D. Representative S-Adenosyl-L-homocysteine (SAH) standard curve used to convert luminescence units to SAH concentration.

E. Sequence of DNA used to wrap di-nucleosomes and 195bp mono-nucleosomes (underlined). Sequence comprises two Widom derived strong positioning sequences with 30bp linker. Linker is engineered for optimum DNMT3 catalytic activity with flanking sites marked in green box and methylated CpG highlighted green.

F. Native gels and SDS-PAGE gels of nucleosomes wrapped with asymmetric 195bp DNA fragment, used in Fig 3B.

G. (left)Native gel showing optimisation of di-nucleosome wrapping by using different ratios of octamer to DNA in the salt gradient dialysis. (right) stability check to see the effect of adding high concentrations of S-adenosyl methionine (SAM) to di-nucleosomes.

H. Formation of DNMT3A1-DNMT3L: di-nucleosome H3Kc36me2 complex. Dose response movement of DNA signal into the wells suggesting di-nucleosomes are bound by DNMT3A1-DNMT3L. Gel resolution was too poor to fully resolve large complex from the wells.

I. Representative cryo-EM micrograph of DNMT3A1-DNMT3L: di-nucleosome H3Kc36me2 complex, with magnified view of boxed particles shown on right. Two adjacent nucleosomes could be observed at a greater frequency than expected given the sample concentration.

**Supplemental Figure S4. Characterising DNMT3A1 recruitment to Polycomb regions via H2AK119ub binding.**

A. EMSA comparing binding of DNMT3A^1-427^ to nucleosomes with or without tri-methylation at H3 Lys27 (H3K27me3). Limiting amounts (2.3 nM) of H3K36me2 nucleosomes were incubated with increasing concentrations (0-8000 nM) of His-MBP-DNMT3A1 constructs. Complexes were resolved by native-PAGE and imaged using blue light excitation and 532nm emission filters. Gels show concentrations 62nM-8000 nM for clarity, quantification was done with full concentration series in duplicate.

B. Methyltransferase assay comparing activity of DNMT3A1-DNMT3L on nucleosomes with nucleosomes modified with H3K36me2 and H3Kc27me3. Methyltransferase activity was detected using Promega MTase-Glo™ Methyltransferase Assay. Experiment done in duplicate. Michaelis-Menten curves were fit using GraphPad Prism 10.

C. Native gels and SDS-PAGE gels of unmodified and modified nucleosomes used in Fig S4 and Fig 4. Denatured gels show equal loading of all four histones and reduced mobility of ubiquitylated H2A. Native gels show shift in mobility of DNA when wrapped into a nucleosome. H2AK119ub nucleosome appears as two bands on gel due to conformation flexibility of the ubiquitin altering mobility.

D. EMSA comparing binding of DNMT3A1 constructs with different lengths of the N-terminal region to H2AKc119ub nucleosomes wrapped with 5’ FAM labelled 175bp Widom601 DNA. Limiting amounts (2.3 nM) of H3K36me2 nucleosomes were incubated with increasing concentrations (0-8000 nM) of His-MBP-DNMT3A1 constructs. Complexes were resolved by native-PAGE and imaged using blue light excitation and 532nm emission filters. Gels show concentrations 25 nM-8000 nM for clarity, quantification was done with full concentration series in triplicate.

E. SDS-PAGE gel of purified minimal UDR and extended UDR DNMT3A1 proteins used in Fig 4E

**Supplemental Figure S5. Cryo-EM data processing scheme for DNMT3A1-DNMT3L:nucleosome H2AK119ub.**

**Supplemental Figure S6. Cryo-EM structure determination and validation for DNMT3A1-DNMT3L:nucleosome H2AK119ub complex.**

A. Representative micrograph of cryo-EM of DNMT3A1-DNMT3L-StrepII on H2AKc119ub nucleosomes.

B. Representative 2D class images from first round of 2D classification in RELION of DNMT3A1-DNMT3L-StrepII on H2AKc119ub nucleosomes. Top views and side views shown, with additional, delocalised non-nucleosomal density seen in the projections.

C. Gold standard Fourier shell correlation (GS-FSC) of the reconstruction of DNMT3A1-DNMT3A1 N-terminal region contains DNA and nucleosome DNMT3L-StrepII determined in RELION. Resolution determined to be 5.1 Å using the 0.143 threshold

D. Euler Angular distribution plot of DNMT3A1-DNMT3L-StrepII:H2AKc119ub nucleosomes. Rod heights and colours are proportional to the number of particles in each direction.

E. GS-FSC curve for final masked map DNMT3A1^UDR^:nucleosome^H2AK19ub^, including unmasked and masked curves. Focused on nucleosome and adjacent density to reach higher detail than above. The dotted line corresponds to 0.143 threshold.

F. Three dimensional FSC for final DNMT3A1^UDR^:nucleosome^H2AK19ub^ map showing global FSC curve (red) and overlap of histogram of directional FSC with the major peak correlating with the global resolution estimate.

G. Euler angle distribution plot of all particles used in the final map. Despite some preferred orientations (red on heat map) no anisotropy in model was observed.

H. Representative regions of the DNMT3A1^UDR^:nucleosome^H2AK19ub^ cryo-EM density map for the different components of the complex. The densities for histones, DNA, and DNMT3A1 UDR region are depicted at a contour level of 0.25 (0.3 DNA) using a final map sharpened map with a b-factor of −50, and the corresponding structural models coloured as in Fig 4B.

**Supplemental Figure S7. Validation of DNMT3A1 UDR nucleosome interaction**

A. Magnified view of DNMT3A1^UDR^:nucleosome^H2AK19ub^ model focusing on the acidic patch interaction and H3-H3’-H2A interaction. Arg-181 interacts directly with carboxylate groups of Glu61, Asp90, and Glu92 in H2A. Arg-167, Arg-169 and Arg-171 form stabilising additional interactions along the acidic patch. Phe-190 and Ala-192 interact in hydrophobic regions, boxed.

B. Native gels and SDS-PAGE gels of modified and unmodified nucleosomes wrapped with 5’ FAM labelled 175bp Widom601 DNA used in Fig5 and Fig S7. H2A.Z runs lower than canonical H2A when unmodified but higher when ubiquitylated in denatured gel.

C. Pull down assay to investigate the effect of DNMT3A1 N-terminal region missense mutations in found in clinical patients on binding to H2AKc119ub nucleosomes. Equal amounts of wild type and mutant His-MBP-DNMT3A1^1-427^ was immobilised on amylose beads and incubated with nucleosomes prior to washing and detection by western blot.

D. SDS-PAGE gel of deletion mutant purified proteins used in Fig 5E.

E. EMSA comparing binding of DNMT3A1^1-427^ to unmodified, H2AKc119ub, H2AZ.1 and H2AZ.1Kc120ub nucleosomes wrapped with 5’ FAM labelled 175bp Widom601 DNA. Limiting amounts (2.3 nM) of nucleosomes were incubated with increasing concentrations (0-8000 nM, 2x dilution series) of His-MBP-DNMT3A1^1-427^. Complexes were resolved by native-PAGE and imaged using blue light excitation and 532nm emission filters. Gels show concentrations 25 nM-8000 nM, quantification was done with full concentration series of two replicates.

**Supplemental Figure S8. Investigating DNMT3A1 ubiquitin site specificity**

A. Native gels and SDS-PAGE showing various ubiquitylated nucleosomes used in DNMT3A1 interaction assays (Fig 6B).

B. EMSA comparing DNMT3A1^1-427^ to unmodified and H2AKc119ub nucleosomes wrapped with 5’ FAM labelled 145, 175 and 195 bp Widom601 DNA. Limiting amounts (2.3 nM) of nucleosomes were incubated with increasing concentrations (0-8000 nM, 2x dilution series) of His-MBP-DNMT3A1^1-427^. Complexes were resolved by native-PAGE and imaged using blue light excitation and 532nm emission filters. Gels show concentrations 7.8nM-8000 nM.

**Supplemental Figure S9. DNMT3A1 UDR region alone does not interact with ubiquitin.**

A. Size exclusion chromatography of DNMT3A1^1-277^ with and without 6xHis-ubiquitin. DNMT3A1^1-277^ (20 μg) was loaded on a Superdex 200 Increase 3.2/300 (Cytiva) column with (grey) or without (black) first incubating with 5x molar excess of 6xHis-ubiquitin (20 μg). An SDS-PAGE gel was run of the fractions of each peak.

B. HSQC NMR of ^15^N ubiquitin with and without DMT3A^164-219^. Spectra were taken of ^15^N ubiquitin (green), then DNMT3A1^164-219^ was added (purple). (bottom) magnified view of assigned residues, highlighting crosspeaks for canonical hydrophobic patch residues (left) and C-terminal hydrophobic patch region (right).

C. Pull down assay measuring binding of 6xHis-ubiquitin to His-MBP-DNMTA1^1-427^. His-MBP-DNMTA1^1-427^ (50 µg) was immobilised on amylose beads (50 μL) and 6xHis-ubiquitin (50 μg) was added. Bound proteins were detected using stainfree UV detection of Mini-PROTEAN TGX Stain-Free Precast Gels. 6xHis-ubiquitin was detected using western blot followed by detection with antibodies against ubiquitin.

D. Pull-down assay using partially purified His-MBP, His-MBP-DNMT3A1^1-427^ and His-MBP-DNMT3A1^1-427^ alanine scanning mutants in the proposed ubiquitin interacting region of the UDR. Proteins were expressed and purified on Nickel affinity beads, eluted and bound to amylose beads prior to incubation with unmodified nucleosomes (left) and H2AKc119ub nucleosomes (right), prior to washing and detection by western blot.

**Supplemental Figure S10. DNMT3A1 activity differs on different nucleosome substrates.**

A. Native gels and SDS-PAGE gels of nucleosomes used for enzymology experiments in Fig 7.

B. EMSA comparing binding of full-length DNMT3A1-DNMT3L-StrepII to 195bp-wrapped nucleosomes used for enzymology experiments (Fig 5C). Limiting amounts (8 nM) of nucleosomes were incubated with increasing concentrations (0-2666 nM, 2x dilution series) of His-MBP-DNMT3A1^1-427^. Complexes were resolved by native-PAGE, stained with diamond stain and imaged using blue light. Gels show concentrations 41nM-2666 nM.

C. Methyltransferase activity of full-length DNMT3A1-DNMT3L-StrepII on unmodified, H3K36me2, H2AKc119ub wrapped with 207bp Widom601 DNA and free 207bp DNA. DNMT3A1-DNMT3L was incubated with increasing concentrations of nucleosomes for 1 hour at 37°C. Methyltransferase activity was detected using Promega MTase-Glo™ Methyltransferase Assay. Experiment done in duplicate. Michaelis-Menten curves were fit using GraphPad Prism 10.

D. Methyltransferase activity of full-length DNMT3A1-DNMT3L-StrepII on unmodified, H3K36me2, H2AKc119ub wrapped with 193bp Widom603 DNA and free 193bp DNA. DNMT3A1-DNMT3L was incubated with increasing concentrations of nucleosomes for 1 hour at 37°C. Methyltransferase activity was detected using Promega MTase-Glo™ Methyltransferase Assay. Experiment done in duplicate. Michaelis-Menten curves were fit using GraphPad Prism 10.

E. Analytical size exclusion chromatography of full-length DNMT3A2-DNMT3L-StrepII. 40 μg protein at 2 mg/ml was loaded on a Superdex 200 Increase 3.2/300 (Cytiva) column. An SDS-PAGE gels was run of all fractions across the peaks.

F. Shift assay comparing DNMT3A1^1-427^ interaction with unmodified, H3K36me2 and Methyl-lysine analogue H3Kc36me2 nucleosomes. Methyl-lysine analogues bind less well compared to true native methylated bond due to differences in the gamma-position on the modified sidechain.

