## Supplementary figures and images for "The N-terminal region of DNMT3A combines multiple chromatin reading motifs to guide recruitment"

### Supplemental Figures

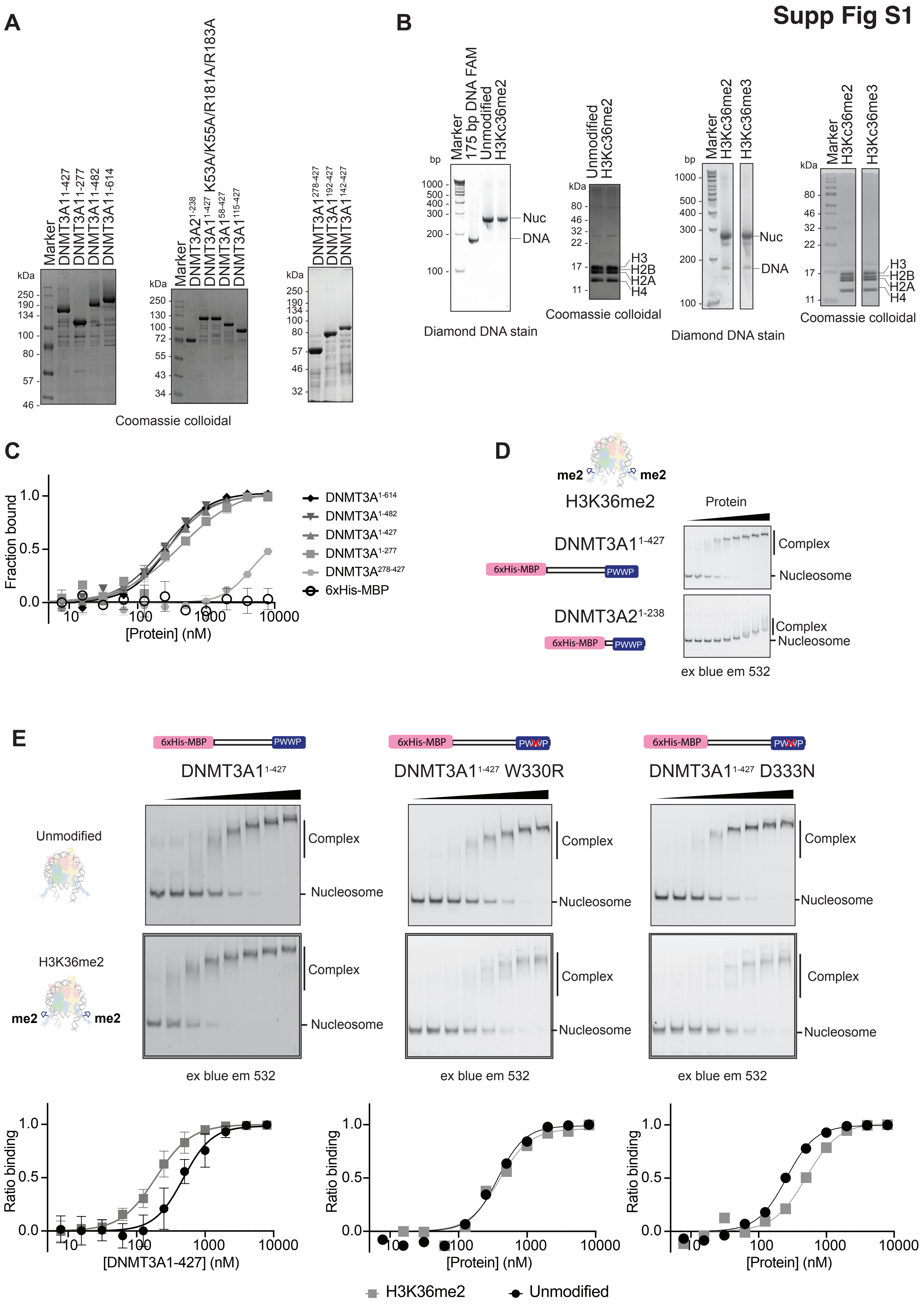

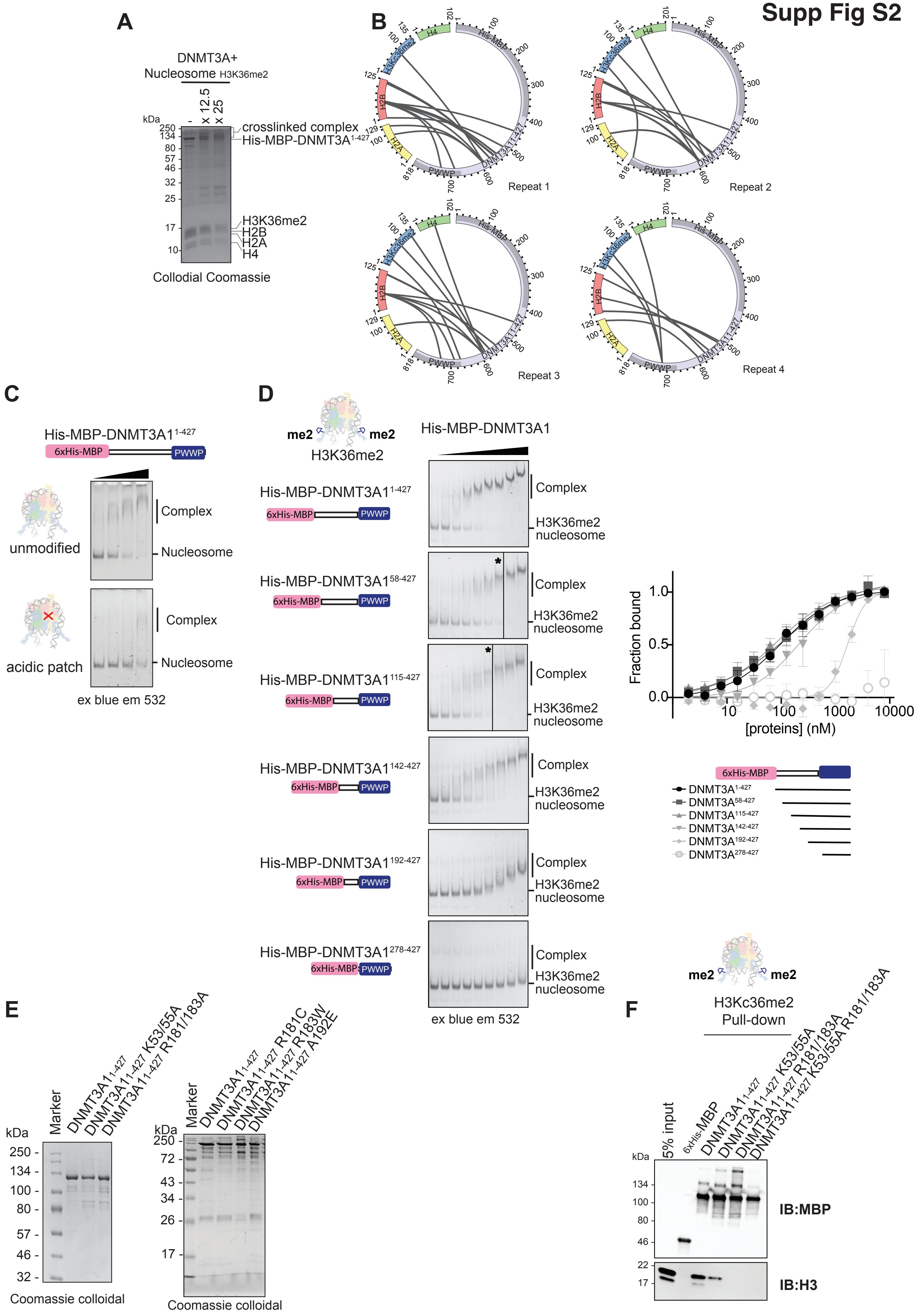

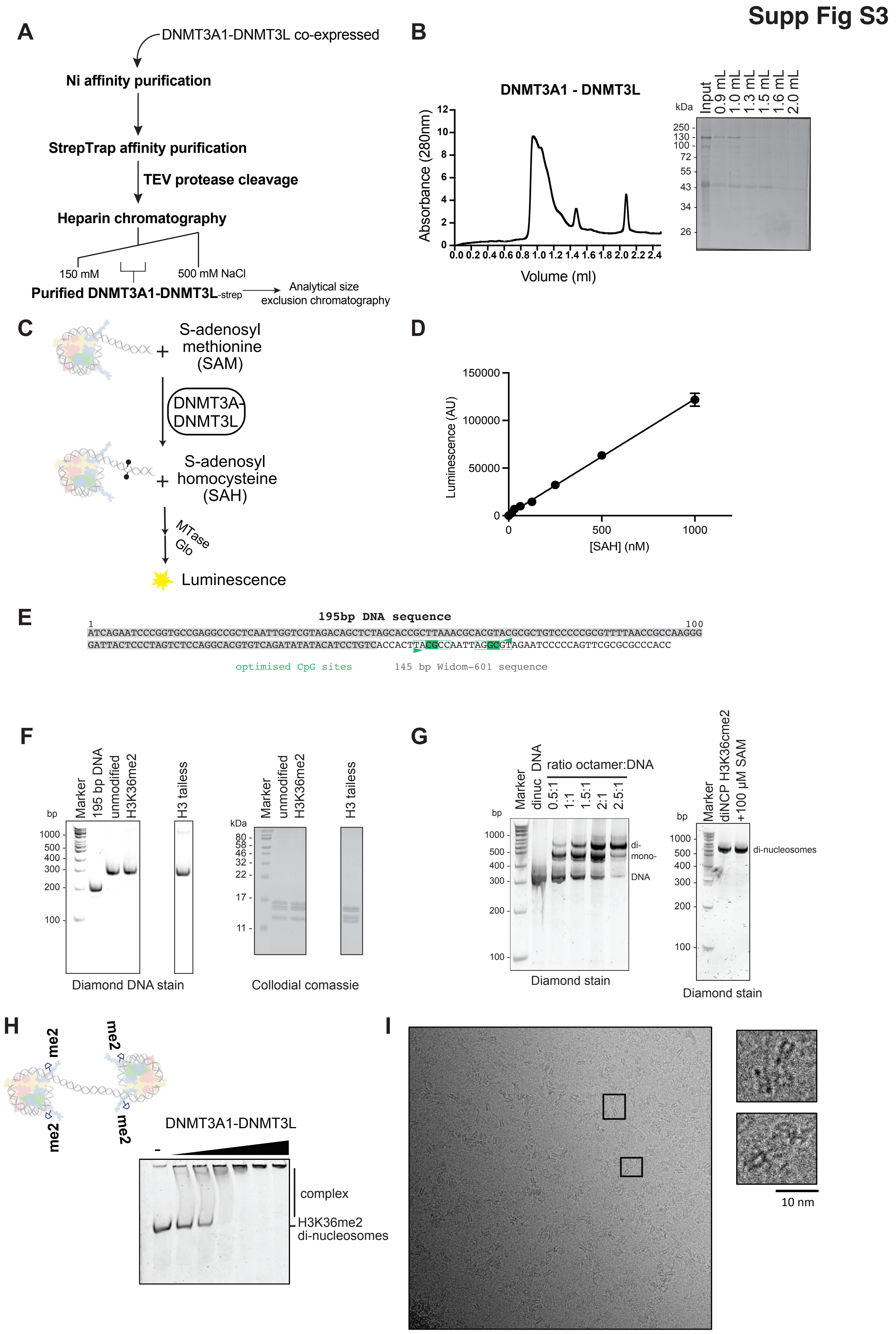

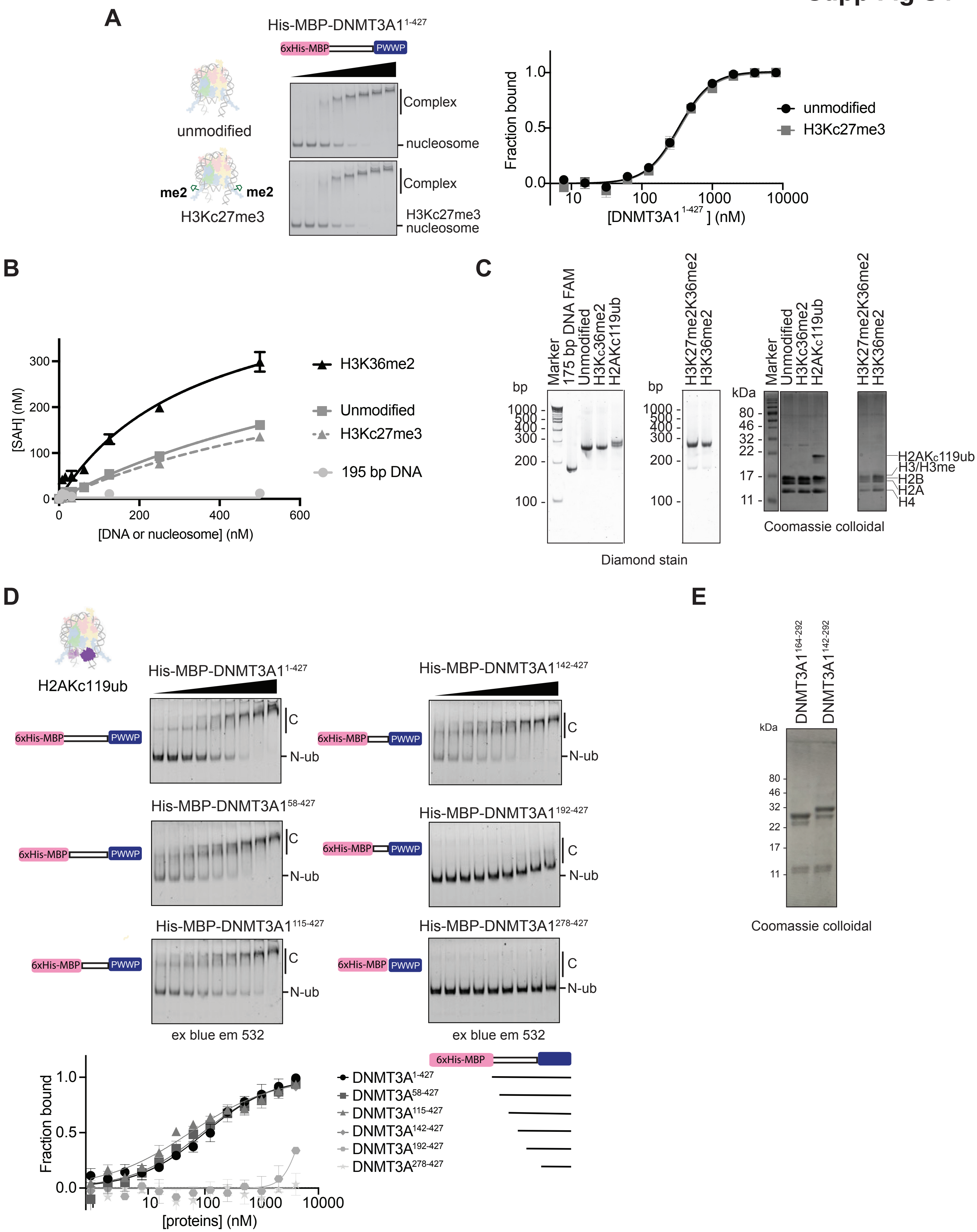

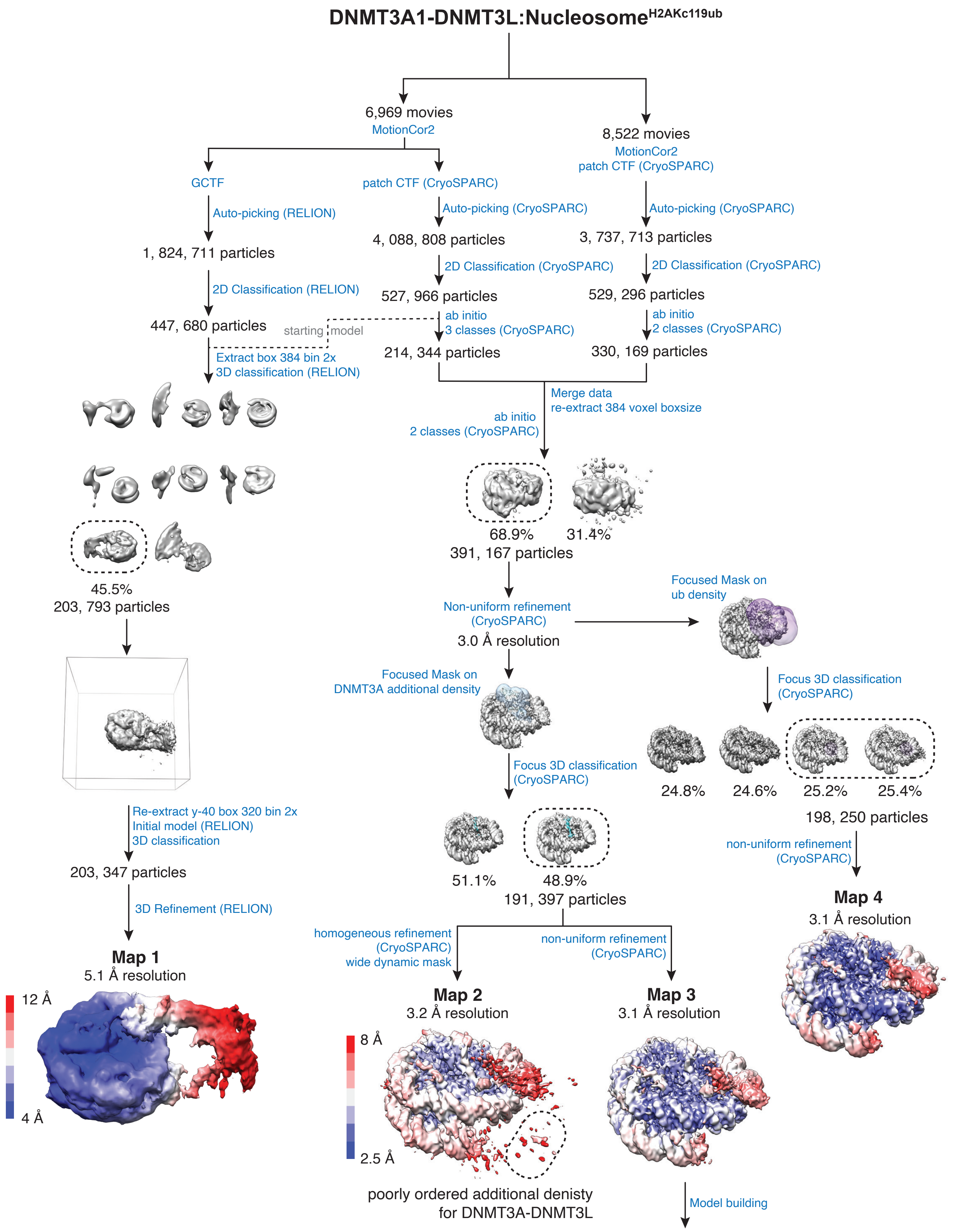

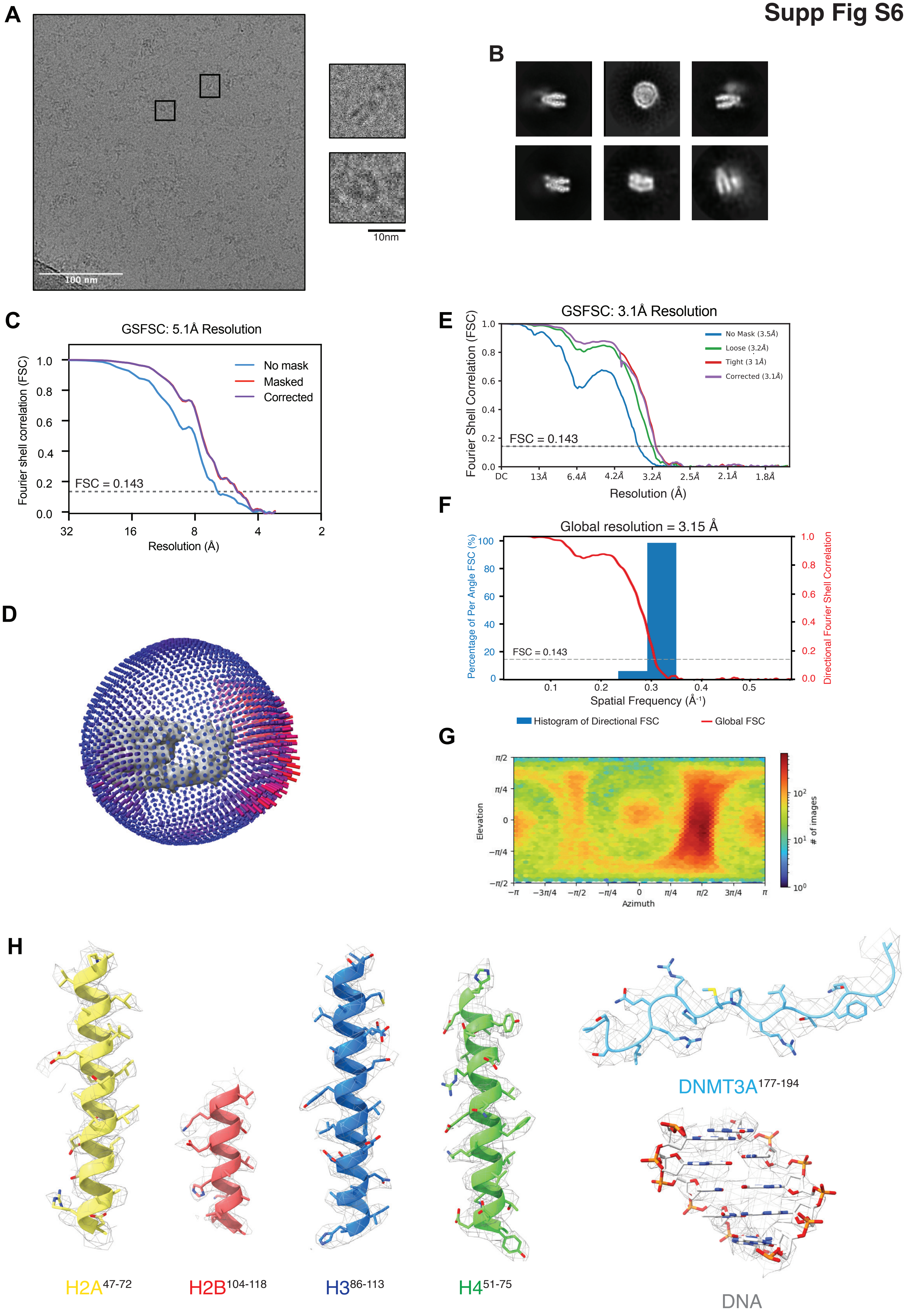

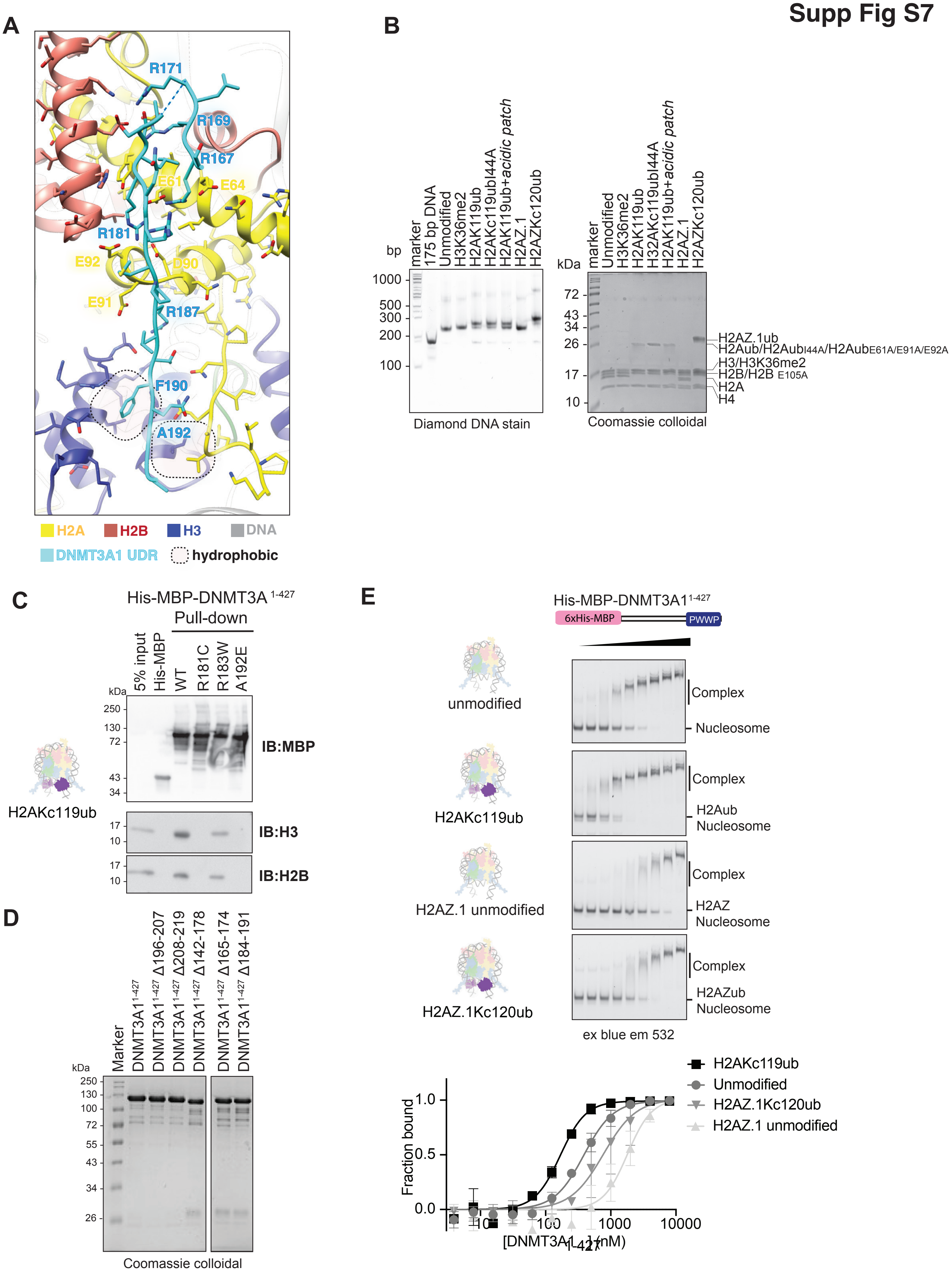

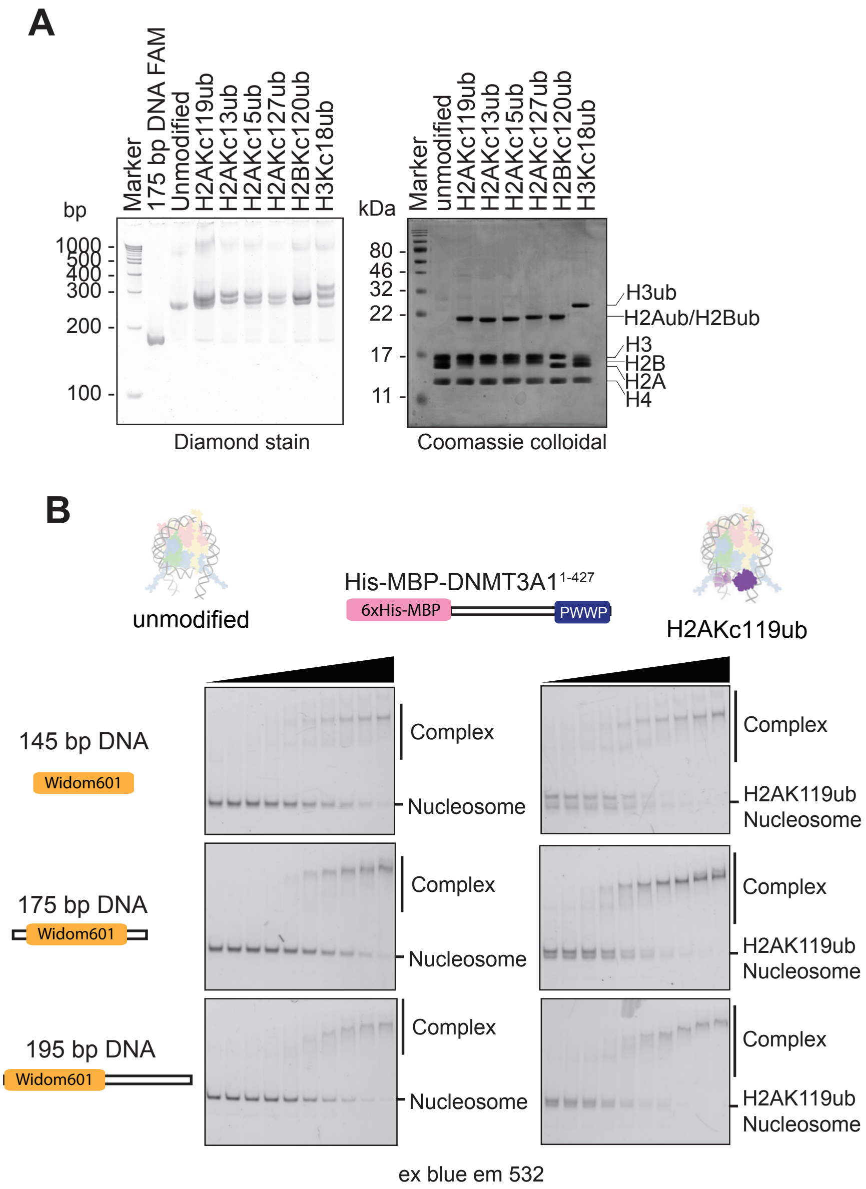

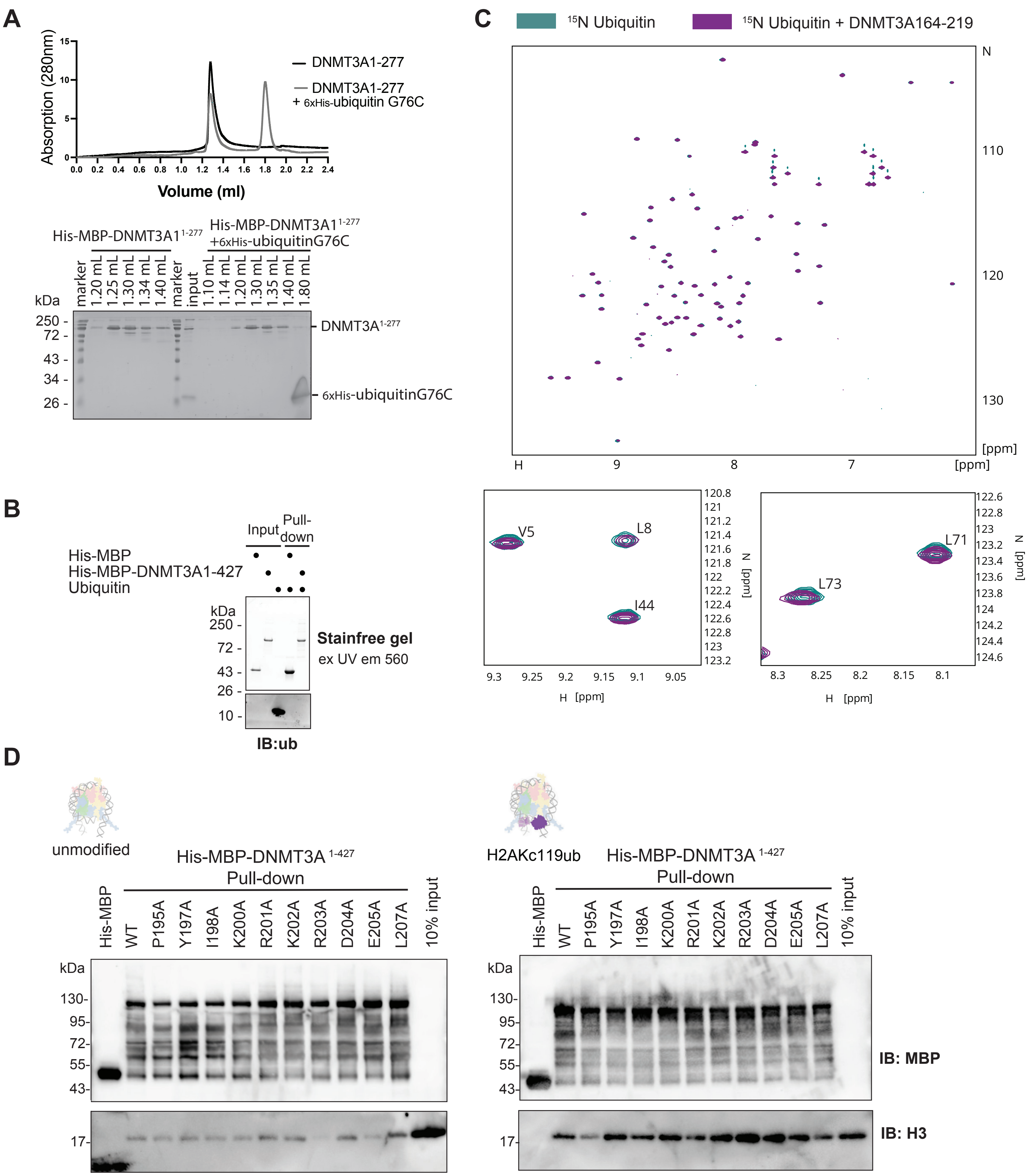

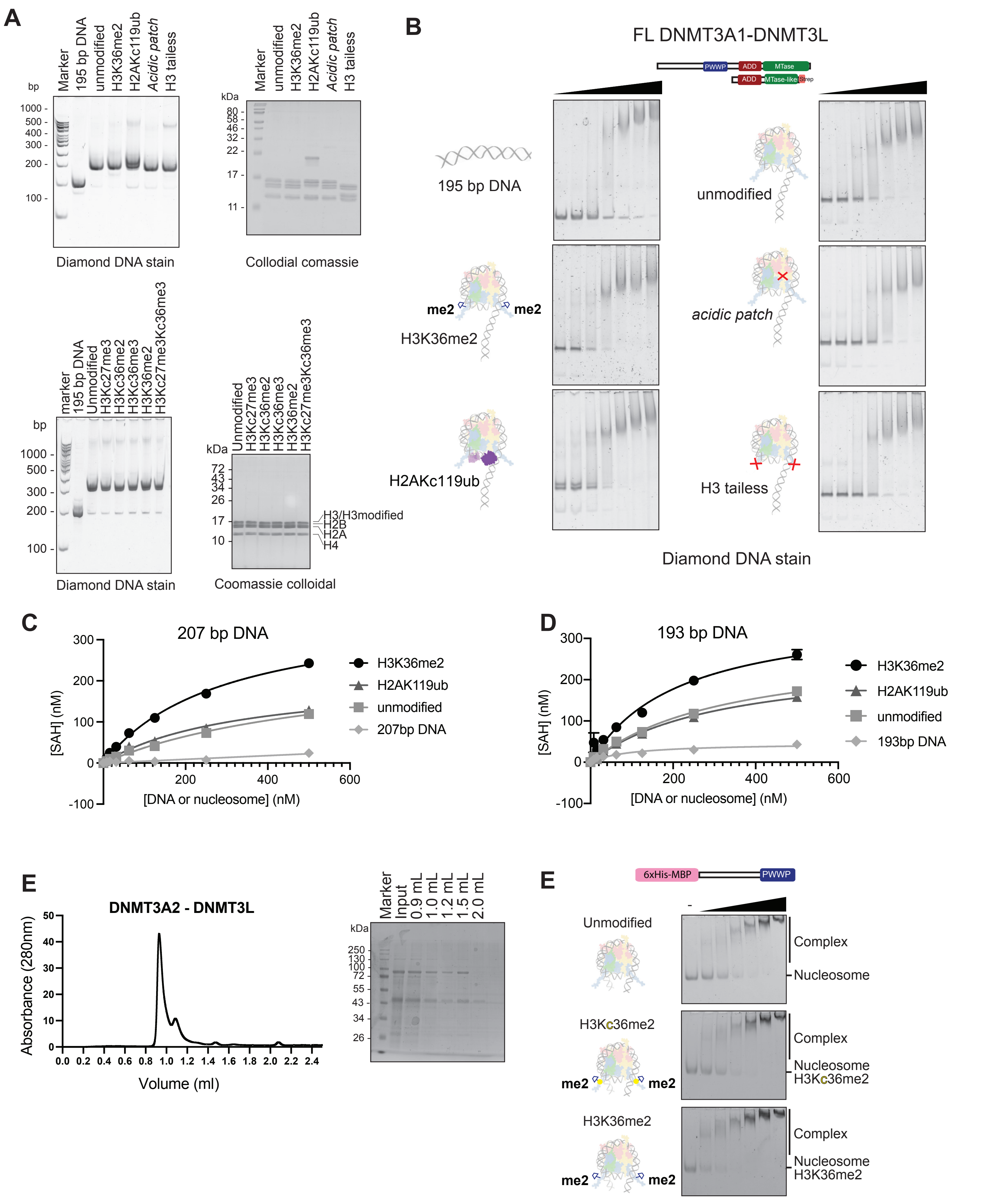
